# The origin of the Gravettians: genomic evidence from a 36,000-year-old Eastern European

**DOI:** 10.1101/685404

**Authors:** E. Andrew Bennett, Sandrine Prat, Stéphane Péan, Laurent Crépin, Alexandr Yanevich, Simon Puaud, Thierry Grange, Eva-Maria Geigl

## Abstract

The Gravettian technocomplex was present in Europe from more than 30,000 years ago until the Last Glacial Maximum, but the source of this industry and the people who manufactured it remain unsettled. We use genome-wide analysis of a ~36,000-year-old Eastern European individual (BuranKaya3A) from Buran-Kaya III in Crimea, the earliest documented occurrence of the Gravettian, to investigate relationships between population structures of Upper Palaeolithic Europe and the origin and spread of the culture. We show BuranKaya3A to be genetically close to both contemporary occupants of the Eastern European plain and the producers of the classical Gravettian of Central Europe 6,000 years later. These results support an Eastern European origin of an Early Gravettian industry practiced by members of a distinct population, who contributed ancestry to individuals from much later Gravettian sites to the west.

## Introduction

Since the 19^th^ century, archaeologists have defined similarities identified across material remains as an archaeological culture, complex, or in some cases, a “people”, and used these definitions to trace the mobility, interactions, and technical development of past populations. The burgeoning field of ancient DNA analysis allows past populations to be studied directly, and the genetic relationships of the manufacturers of materially defined cultures can now be characterized. While palaeogenomics and archaeological culture describe two separate past phenomena, they are not always unrelated, and Upper Palaeolithic (UP) Europe has shown surprising correlations between closely-related genetic clusters and archeologically defined material industries^1^. The Gravettian technocomplex defines European Mid-Upper Palaeolithic (MUP) industries characterized by a suite of shared innovations, along with specific modalities of their production, such as stone Gravette points, backed blades and bladelets, personal ivory and shell ornaments, ochre, and antler or bone tools. The Gravettian became widespread throughout Europe beginning after ca. 36,000^2^ until ca. 23,000^3^ cal BP years ago (ca. 32-21 ka ^14^C BP). The technocomplex defined by this term, however, is by no means homogeneous, describing rather shared practices across many regional *facies* and evolving *stages*^3^ (See Supplementary Text). While the Gravettian is considered to be a local European industry, its origins continue to be the subject of ongoing research and debate^4^. The density of Gravettian sites found in the Danubian valley radiocarbon dated as early as ca. 36,000 years cal BP have been used to argue for a local evolution of the Gravettian from the technical legacy of the Aurignacian in the upper valley^5^, or through acculturation with post-Mousterian leaf-point transition industries, such as the Szeletian, in the Middle Danubian basin^6^. However, long-recognized similarities between features of Gravettian lithic traditions and Near Eastern industries, such as the Ahmarian, which is found 10,000 years earlier in the Levant, as well as similar Early-Upper Palaeolithic (EUP) micro-laminar industries found in both northern and southern slopes of the Caucasus suggest an earlier influence from the southeast^7–9^.

Although the integration of Eastern European EUP and MUP traditions into an archaeological framework defined in Central and Western Europe has historically been problematic, recent reassessments of EUP assemblages from the Caucasus^10–12^ emphasize similar lithic traditions among roughly contemporaneous (40-30,000 cal BP) layers at sites such as Dzudzuana Cave (layer D), Ortvale Klde (layer 4c), Mezmaiskaya Cave (layer 1C), and the six layers of Buran Kaya III (layer 6-5 to 5-2) in Crimea^13,14^ (see Supplementary Text for the stratigraphy of Buran-Kaya III). Some authors suppose that the appearance of these Caucasian and Eastern European EUP assemblages, which include backed blades and bladelets, are primarily based on the distribution and transformation of Ahmarian traditions from Near East to Europe, probably through the Caucasus, independent of that which brought the proto-Aurignacian to Central and Southern Europe and the Mediterranean area^10,14–16^. The parallels between these Eastern European industries and the Gravettian appearing later in Central Europe and the Danubian Valley have led some to propose the term “Early Gravettian” to describe these industries to distinguish them from the classical Gravettian^8,16,17^. However, the relationship of this Early Gravettian identified in Eastern Europe to the development of the Gravettian in the west, as well as the extent to which it may have involved transfers and adaptations of technology and movements of people, are unclear. The direct study of the populations associated with these industries through genomic analysis of skeletal remains from key archaeological sites will likely clarify this process and answer important questions concerning how technologies disperse and change in UP societies.

Several human remains associated with the classical Gravettian from Central Europe, Belgium and Italy were previously analysed genetically and found to share more genetic drift with each other than individuals associated with other material cultures^1^. This group was termed the *Vestonice cluster*, after Vestonice16, the best-covered genome from the ca. 30,000-year-old (cal BP) Gravettian site Dolní Věstonice II in the Czech Republic. Interestingly, Vestonice16 and other high-coverage genomes from the Gravettian sites Krems-Wachtberg (Austria) and Ostuni (Italy) have been found to be admixed, deriving between 40 to 90% of their ancestry from a lineage related to Kostenki14^1,18^, an individual living ca. 7,000 years earlier at the Kostenki-14 site found in the Borshchyovo archaeological complex of Western Russia, linking Gravettians of Central Europe and the Apennine peninsula with an earlier population farther east.

While the Kostenki14 burial has no associated cultural material, contemporaneous Gravettian artefacts have been recovered in Eastern Europe from three cultural layers at Buran-Kaya III in Crimea and dated between ca. 38-34,000 cal BP^2,19^. Discovered in 1990, the Buran-Kaya III rock shelter contains stratigraphic layers spanning from the Middle Palaeolithic to the Middle Ages. The Gravettian layers have yielded backed microliths, microgravette points, ochre, body ornaments of ivory, shell, and teeth, as well as multiple human skull fragments showing signs of post-mortem processing^2^. Buran-Kaya III is the earliest site bearing Gravettian material^20^ (see Supplementary Text), yet it remains a distant outlier, both geographically and temporally, to the more densely clustered appearances of the Gravettian occurring several thousand years later and 2,000 km to the west in the Danubian valley and the Swabian Jura, such as Krems-Hundssteig and Geiβenklösterle, respectively.

To genetically characterize one of the earliest manufacturers of the Gravettian complex and how they relate to other known MUP populations, as well as to further explore the relationships between UP populations and the material culture they made, we present genome-wide data from a human parietal bone fragment, BuranKaya3A, aseptically excavated in 2009 from layer 6-1 of Buran-Kaya III (see Supplementary Text). Radiocarbon (AMS) dates from a different human cranial fragment from this layer range from 36,260 to 35,280 cal BP (31,900+210-220 ^14^C BP)^2^ (Figure 1 and Table S1), which is located between two other Gravettian layers, while all dated material (n=4) from the same layer ranges from 37,560-33,850 cal BP^19^.

**Figure 1.**
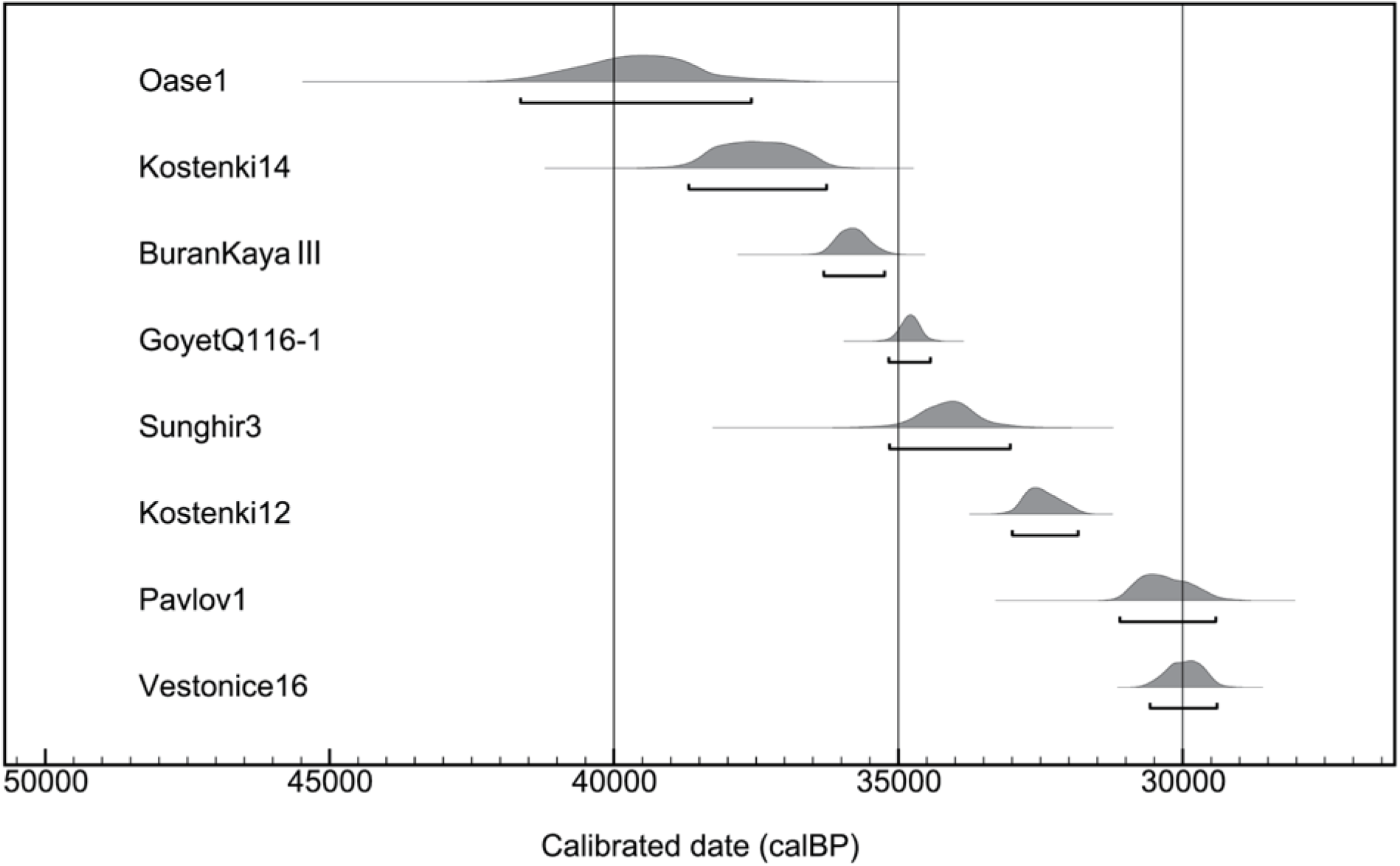
Recalibration of comparative AMS ^14^C dates of Buran-Kaya III, layer 6-1 with Early and Mid-Upper human Palaeolithic European samples. All dates were recalibrated using the software OxCal v4.3.2 based on the IntCal13 calibration data set^38^. For complementary information on cultural contexts, see Table S1.

## Results

Initial genetic characterization of BuranKaya3A revealed extremely poor preservation of endogenous DNA, both in fragment length and in low quantity relative to environmental DNA, but little contamination with modern human DNA as assessed by quantitative PCR. In order to identify the most efficient material for mitochondrial enrichment and shotgun genome sequencing, our strategy was to 1) screen single-stranded DNA libraries by both shallow shotgun sequencing and mitochondrial enrichment from eight extractions taken from four different areas of the sample and subjected to two different extraction treatments, 2) identify the extract that performed best in terms of highest sequence complexity, endogenous content, and lowest modern human contamination, and 3) construct a second series of single-stranded libraries with and without UNG for deeper mitochondrial enrichment and shotgun sequencing. The best candidate identified from the screening results contained 0.34% endogenous DNA (see *Methods* section and Extended Data Figure 1). The second series of libraries from this extract was used to generate 82-fold coverage of the mitochondrial genome and shotgun nuclear data (Table S2).

### Mitochondrial and Y-chromosome haplogroups

The mitochondrial haplogroup of BuranKaya3A was determined to belong to an early branch of the N lineage, N1. Surprisingly, this assignment falls outside of the lineages previously reported for UP Europe, nearly all of which derive from later N branches (U and R haplogroups, Figure 2). The N1 of BuranKaya3A is notably distinct from the mitochondrial haplogroup N identified from the roughly 40,000-year-old mandible from Peștera cu Oase in Romania, which belongs to a more basal branch that has no modern descendants^21^. In addition, the N1 of BuranKaya3A carries three of the eight mutations occurring prior to N1b, a rare haplogroup most highly concentrated in the Near East, yet appearing broadly from western Eurasia to Africa. The descendants of the N1b node include N1b2, currently found only in Somalia^22^, and N1b1b, found in nearly 10% of Ashkenazi Jewish haplogroups^23^. These three mutations allow us to place BuranKaya3A on a lineage apart from that which has been proposed to later enter Europe from Anatolia during the Neolithic (N1a1a)^24^. Among ancient samples, the mitochondrial sequence of an 11,000-year-old Epipalaeolithic Natufian from the Levant (“Natufian9”)^25^ is also a later derivative of this N1b branch. Thus, mitochondrial sequences branching both upstream and downstream of the BuranKaya3A sequence can be traced to the Near East, and the modern presence in Europe of haplogroups descended from the N1 (N1b1b and N1a1a) branch to which BuranKaya3A belongs appear to be due to later migrations from the Near East (Extended Data Figure 2). We determined the genetic sex of BuranKaya3A to be male using both the ratio of chromosome X and Y mapped reads, giving an R_y_ value of 0.0893-0.097 (95% CI, SE 0.002)^26^, as well as a ratio of chromosome X mapped reads to the average of autosomal reads of 0.55 (a ratio near 1.0 would indicate diploid for X). From the reads mapping to the Y chromosome, six out of six Single Nucleotide Polymorphisms (SNPs) that overlap with diagnostic sites for Y-haplogroup BT all carry the derived allele, allowing a minimum assignment to BT, which has origins in Africa, with additional derived alleles suggesting an eventual placement of CT or C, found in Asia and the Epipalaeolithic Near East^25^. Additional ancestral alleles make an assignment of C1a2 or C1b, which appear in UP Europe^1^, unlikely (see Table S3 for a summary and comparative placement of Palaeolithic Y-haplogroups, and Supplementary Data 1 for a complete list of Y diagnostic SNPs).

**Figure 2.**
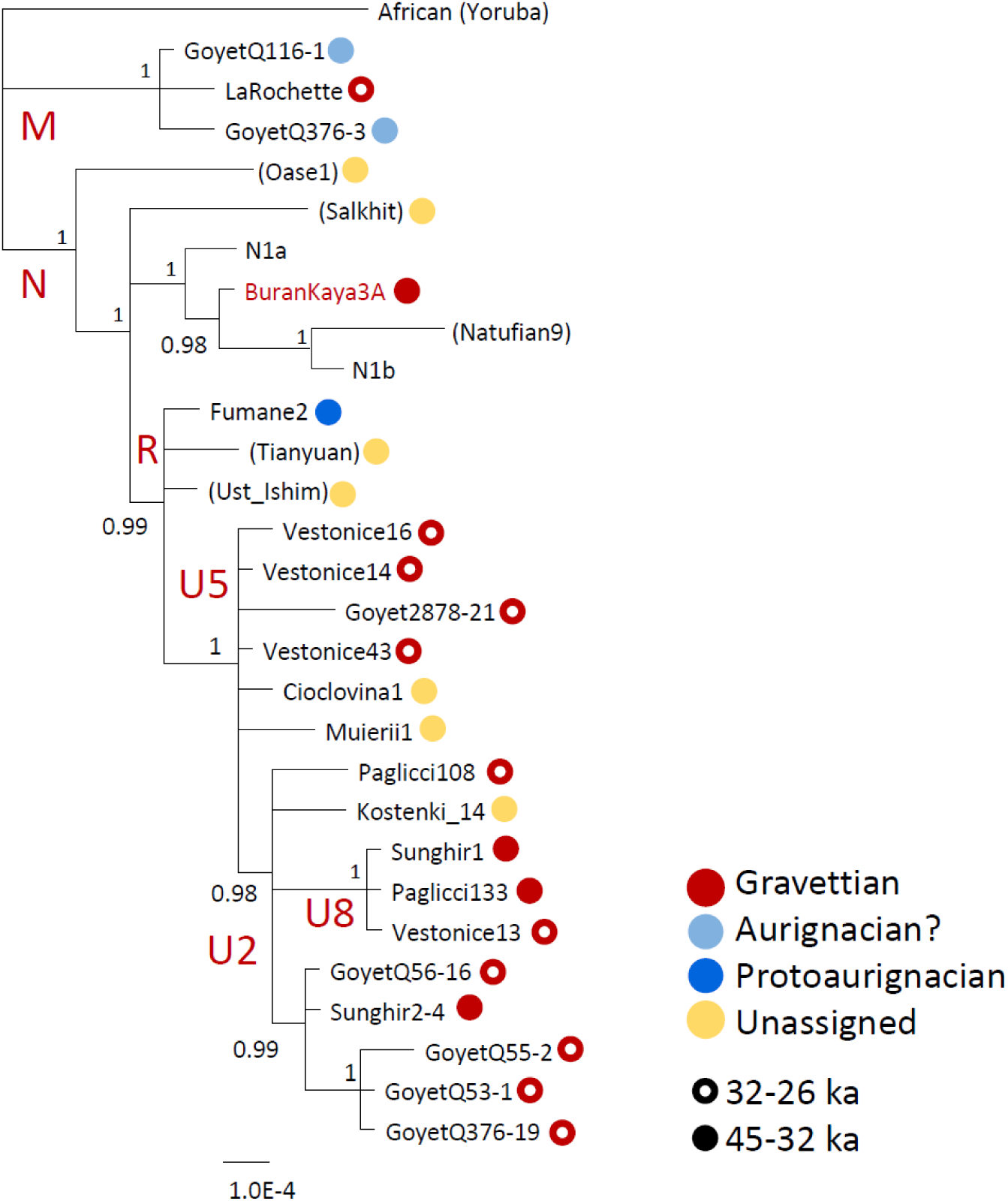
Bayesian phylogenetic tree of mitochondrial sequences, excluding Hyper-Variable Regions, from Early and Mid-Upper Palaeolithic individuals including BuranKaya3A. Posterior probability indicated at the nodes. Non-European individuals are in parentheses. General ages shown as either closed or open circles, and pre-glacial material cultures, when known, indicated by colour. A question mark by the Aurignacian indicates a cultural assignment by dating rather than direct association. Scale bar denotes substitutions per site. Sources for mitochondrial sequences are listed in Table S5.

### Neanderthal ancestry

Neanderthal settlements, attributed to the Micoquian, Kiik-Koba type, at Buran-Kaya III (layer B) has been dated by faunal bone fragments to 43.5 to 39.6 ka cal BP^19^, demonstrating an earlier Neanderthal presence at the site just prior to the Campanian Ignimbrite eruption (39,280±110 years cal BP (^40^Ar/^39^Ar))^27^. Anatomically Modern Human (AMH) remains dating to this period in Romania have documented local admixture with late Neanderthals in Eastern Europe^21^, leading us to investigate whether admixture with local Neanderthals in Crimea could be detected in early AMHs living ca. 4,000 years after the presumed disappearance of Neanderthals from the region^19,28^. Neanderthal ancestry calculated using ancestry informative SNPs^1^ on libraries from four independent preparations determined that BuranKaya3A possesses 3.4% (SD 0.008) Neanderthal ancestry (Table S4). This level of Neanderthal ancestry is typical for contemporaneous West Eurasians and shows no evidence of late, local Neanderthal admixture in Crimea with AMHs ancestral to the population to which BuranKaya3A belonged.

### Genomic relationships of UP Europe and their archaeological cultures

The genetic relationship between BuranKaya3A and other UP individuals for which sufficient genomic data exist can be measured using outgroup *f*3-statistics and various D-statistical tests^29^. These were performed using SNPs from 52 previously published individuals dating from the Initial Upper Palaeolithic (IUP) to the Mesolithic that had been reprocessed in the same pipeline used for BuranKaya3A to reduce possible artefacts that may arise from data coming from disparate sources. 27,774 SNPs having base quality 30 or above were identified in BuranKaya3A that overlapped with the nearly 3M SNPs from the combined panels in Fu *et al*. 2016^1^. To minimize standard errors, we restricted the results of this analysis to include only individuals having more than 300,000 SNPs overlapping with the panel.

Outgroup *f*3-statistics quantify the amount of shared genetic drift between two populations relative to an outgroup population and have been used to reveal genetic affinities among UP individuals^1^. Of the 21 ancient individuals (*x*) including at least 1000 SNPs in the calculation: *f*3(BuranKaya3A, *x*; Mbuti) or *f*3(BuranKaya3A, *x*; Han), the highest *f*3 values for both outgroups were found when *x* was either Sunghir3 (SIII), a 34,000-year-old (cal BP) group burial characterized as either Streletskian^30^ or Eastern Gravettian^31,32^ (see Supplementary Text) found ca. 1,300 km to the northeast of Buran-Kaya III, Vestonice16, from the context of a Gravettian *facies* known as the Pavlovian, who lived ca. 6,000 years later and ca. 1,400 km to the northwest, and Kostenki14, a burial predating BuranKaya3A by ca. 1,000 years and lying ca. 1,100 km to the northeast (Figure 3). Notably, BuranKaya3A shows less affinity for the 35,000-year-old (cal BP) GoyetQ116-1 from Goyet cave in Belgium, with no direct cultural association, but dated to the Aurignacian period in this area^1,33^, as well as the El Miron cluster, which corresponds to European late glacial (Magdalenian) hunter-gatherers^1^. BuranKaya3A was also found to be particularly distant from the Villabruna cluster, representing European post-glacial hunter-gatherers (Figure 3 and Extended Data Figure 3A-B). The individual from Buran-Kaya III can thus be defined as a member of a population making an Early Gravettian in Eastern Europe whose closest known genetic relationships are with populations living within a 3,000-year window (ca. 37,000-34,000 cal BP) farther northeast on the Eastern European plains (Sunghir3 and Kostenki14), as well as with Gravettian populations who appeared ca. 6,000 years later in Central Europe (Vestonice16).

**Figure 3.**
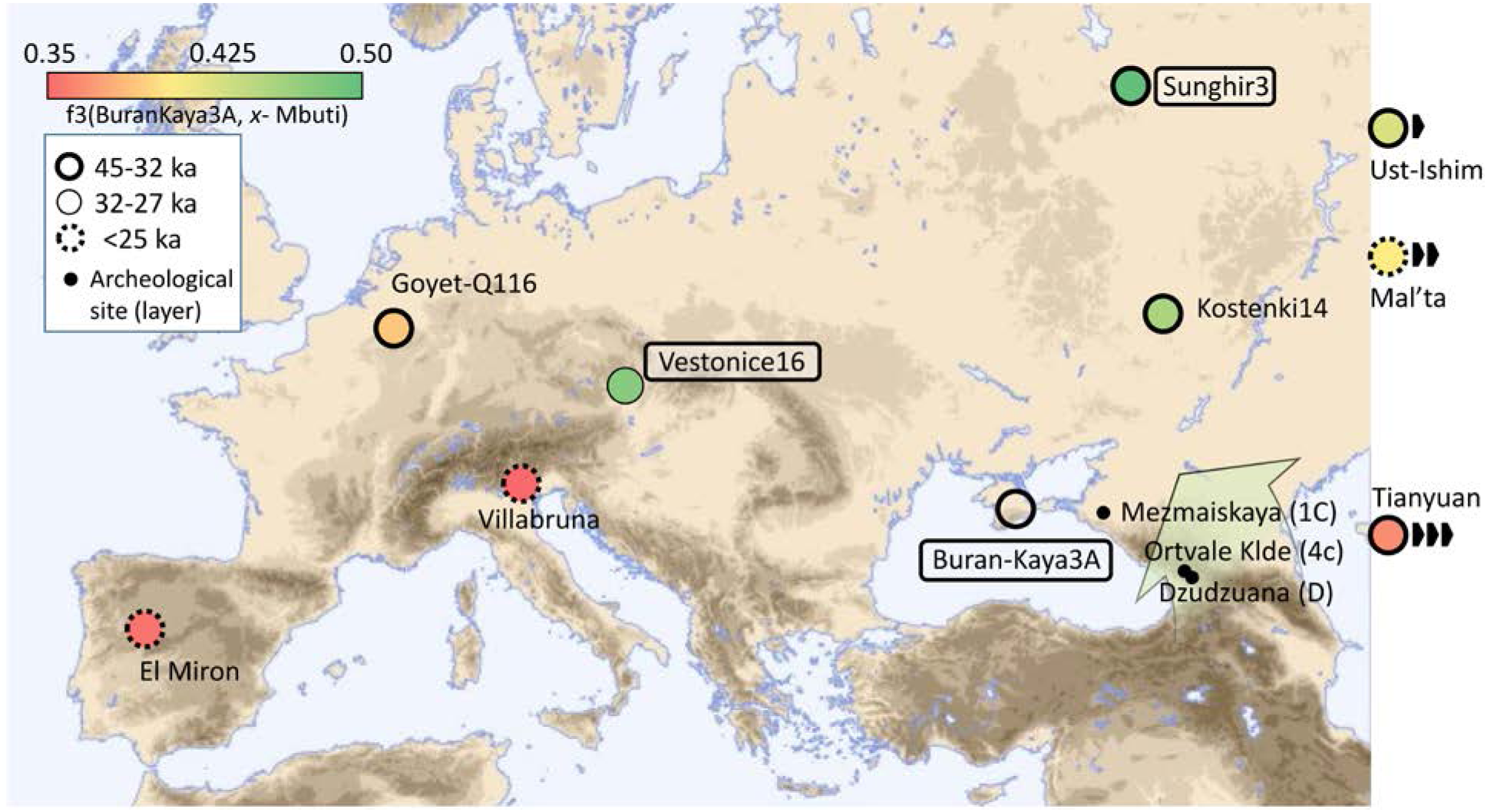
Location and heatmap of *f*3(BuranKaya3A, *x*; Mbuti) with archaeological sites mentioned in the text. Values of high genomic coverage ancient samples (higher values show more shared alleles with BuranKaya3A). Boxed text indicates samples associated with a Gravettian archaeological context (including Sunghir3, alternatively described as Streletskian). Non-European samples are given along the right margin at their approximate latitudes, their relative distances indicated by black arrows. Archaeological sites for which no human genetic data are available that contain micro-laminar industries comparable with Buran-Kaya III at contemporaneous layers (in parentheses) indicated by black dots. A broad green arrow shows the proposed EUP route introducing the Early Gravettian into Europe, suggested by the similar features of these assemblages and the lack of a Common Western Eurasian genomic component in BuranKaya3A.

We then applied D-statistics to measure shared drift between BuranKaya3A and the individuals tested above (*x*) as compared to the 45,000-year-old (cal BP) Central Asian Ust-Ishim (*D*(*x*, Ust-Ishim; BuranKaya3A, Mbuti)), shown to be basal to all western MUP Eurasians studied to date^34^ (Figure 4a). Substituting the modern Han for Ust-Ishim gives additional support for the above relationships with greater significance (Z-scores > 2.5) (Figure 4b). Full results for all tested samples using both all SNPs and transversions only are given in Extended Data Figures 4 and 5. The D-statistics support the results of the outgroup *f*3-statistical tests, except that here Sunghir3 is replaced by Vestonice16 as sharing the most alleles with BuranKaya3A, although we note results from both Vestonice16 and Sunghir3 overlap within one standard error. The additional D-statistic *D*(*w*, *x*; BuranKaya3A, Mbuti), where w and x are various well-covered ancient individuals representing previously defined Eurasian populations, was unable to significantly resolve which of these two individuals is most closely related genetically to BuranKaya3A: the ca. 34,000-year-old (cal BP) East European Sunghir3 or the ca. 30,000-year-old (cal BP) Central European Vestonice16 (Extended Data Figure 6).

**Figure 4.**
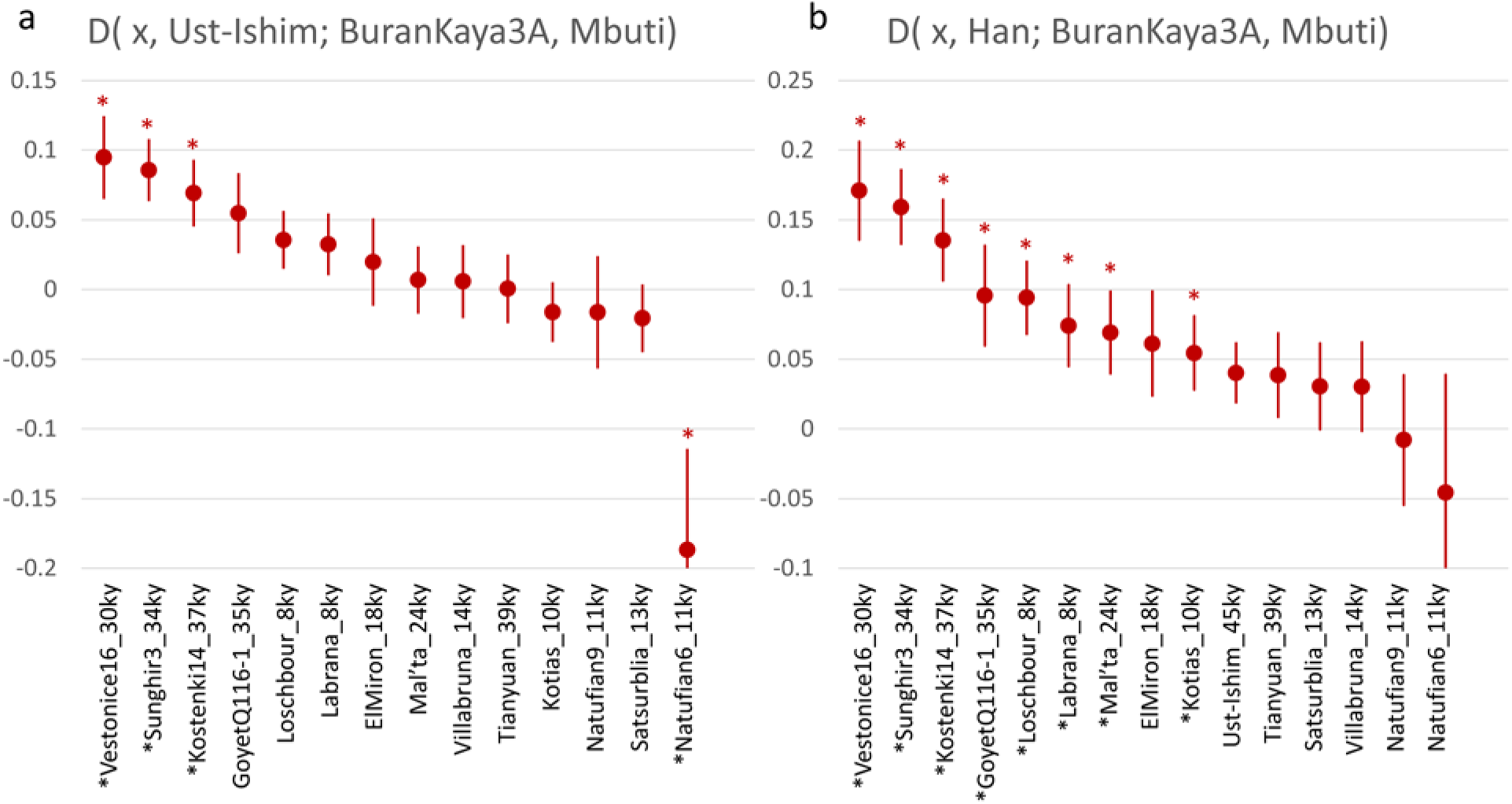
D-statistics results for a) D(*x*, Ust-Ishim; BuranKaya3A, Mbuti) and b) D(*x*, Han; BuranKaya3A, Mbuti) where *x* = selected UP and Mesolithic Eurasians. Starred individuals denote a significant Z-score >2. Approximate ages, being the mean of the latest calibrated published date rounded to the nearest ky, are appended to the names. More positive values represent increased allele sharing between *x* and *BuranKaya3A* relative to *Ust-Ishim* or *Han*, with *Mbuti* as an outgroup. Results from all SNPs shown. All Z-scores and results from transversions only are given in Extended Data Figure 4. The values for Kotias, Satsurblia, and the Natufians are distorted due to Basal Eurasian content in these samples (see Extended Data Figure 9). Error bars = one standard error.

Genomic data from less well covered MUP Europeans allows us to better define the relative position of BuranKaya3A with more individuals from Gravettian contexts, including representatives of two of the three previously identified *Vestonice* genetic sub-clusters corresponding to Gravettian sites^1^. Comparing two low-coverage individuals requires more caution in interpreting the results due to the low number of SNPs used for the calculations. However, we were able to establish additional support for the relationships by applying a strict filtering process whereby 1) we performed the analysis using two datasets, one including all SNPs and the other considering only the transversions. The transversion-only dataset reduces the number of SNPs used, but eliminates statistical noise from deaminated cytosines, a common form of damage in ancient DNA. 2) We excluded samples using less than 50 SNPs for *f*3 analysis in the smaller transversion-only dataset. 3) We required the resulting values from both datasets to be within 30% of each other. This approach allowed us to define genetic affinities between BuranKaya3A and the following individuals: Kostenki12, a 32,500-year-old (cal BP) neonate whose remains were found at the Kostenki-Borshchyovo archaeological complex in Russia and associated with a cultural layer attributed to a local industry defined as the Gorodtsovian^35^; Vestonice43 and Pavlov1, two Central European Gravettian (Pavlovian) burials similar in age and location to Vestonice16 and belonging to the “Vestonice Central European” genetic subcluster; Paglicci133, a ca. 33,000-year-old (cal BP) tooth from the Apulia region of Southern Italy found in association with Early Gravettian cultural material; and Ostuni1, a ca. 27,500 year-old (cal BP) Gravettian burial, also from the Apulia region and both belonging to the “Vestonice Italian” genetic subcluster. No members of the “Vestonice Goyet” subcluster from Goyet cave in Belgium passed our filters to be included in the analysis (full descriptions of these samples and clusters are given in the Supplementary Information of Fu et al. 2015^1^). Visual representation of the outgroup *f*3-statistics, *f*3(BuranKaya3A, *MUP*; Mbuti), for all samples passing filters is shown in Extended Data Figure 7 and full results for all tested samples are given in Extended Data Figure 8.

Of the less well-covered samples, Kostenki12 is shown to have a high affinity with BuranKaya3A. This is in agreement with the previously reported affinity between Kostenki12 and Sunghir3^36^ and closely links BuranKaya3A to all EUP and MUP individuals currently known from the Eastern European plains. To the west, BuranKaya3A shows close genetic affinity with Pavlov1, followed more distantly by Vestonice43 and Ostuni1, and at a greater distance, the older Paglicci133. In summary, these results allow us to firmly establish a close genetic relationship between BuranKaya3A with contemporary EUP and MUP populations of the Eastern European plains, as well as the broader population manufacturing the Pavlovian *facies* of the Gravettian in Central Europe ca. 6,000 years later, but less so with those associated with the Gravettian of southern Italy. This implies a network of gene flow across the Eastern European plains and the Danubian Valley between 37,000-30,000 years ago (cal BP) which, apart from the local Gorodtsovian culture found at Kostenki 12, can be associated with various *facies* of the Gravettian technocomplex. The results from Ostuni1 and Paglicci133 indicate a more indirect relationship between these populations and BuranKaya3A. Higher coverage genetic information from early Gravettian sites in Italy will be required to better understand to what extent the movements of people and ideas lead to the appearance of the Gravettian in the southern Apennine Peninsula.

### Low Common West Eurasian ancestry in UP Eastern Europe

Ancestry from a population which split from all non-Africans prior to their separation from each other, termed Basal Eurasian, had not been known in Europe until after the Last Glacial Maximum (LGM)^1^. However, two 24,000-27,000-year-old (cal BP) individuals from layer C at Dzudzuana Cave in the southern Caucasus have recently been reported to share ~70% common ancestry with Villabruna and ~30% ancestry deriving from this Basal-Eurasian source^18^. Additionally, varying layers of Villabruna ancestry, which did not enter Europe in an unmixed form prior to ca. 14,000 years ago (cal BP)^1^, have been found in members of the Gravettian Vestonice cluster as well as the Magdalenian El Miron cluster, indicating a degree of shared ancestry in these groups with a “Common West Eurasian” population^18^. We calculated the relative level of Basal Eurasian ancestry using D-statistic *D*(*EUP East Asian*, *UP*, Ust-Ishim, Mbuti), where a positive value indicates Basal Eurasian ancestry as allele sharing between the UP individual and Africans. This analysis showed BuranKaya3A, like other pre-LGM UP Europeans, to be lacking Basal Eurasian ancestry (Extended Data Figure 9). Similar tests examine the relative affinities of Palaeolithic and Mesolithic populations to Villabruna as opposed to either modern East Asians, *D*(*x*, Han; Villabruna, Mbuti), or BuranKaya3A, *D*(*x*, BuranKaya3A; Villabruna, Mbuti) (Extended Data Figures 10 and 11, respectively). These results show levels of Villabruna ancestry in Eastern European EUP individuals (Sunghir3, Kostenki12, Kostenki14, and BuranKaya3A) below that of later UP Central and Western Europeans from Gravettian contexts (the Vestonice cluster). Intriguingly, this includes a proportion of shared ancestry with Villabruna in BuranKaya3A similar to that found in Ust-Ishim and both ancient and modern East Asians (Extended Data Figures 10 and 11), which should be insignificant based on previous admixturegraph analysis^18^.

## Discussion

The results of the genome-wide analysis of BuranKaya3A offer important evidence linking the previously established genetic signature of the manufacturers of the Gravettian in Central Europe to a much earlier appearance of the Gravettian in Eastern Europe. The absence of the “Common West Eurasian” ancestry, as represented by Villabruna, in BuranKaya3A marks a key genetic distinction between the Gravettian inhabitants of Buran-Kaya III, possibly including the broader populations of EUP Eastern Europe as well, and the UP populations of Western and Central Europe, which is characterized by a West-to-East reduction in “Common West Eurasian” ancestry (seen in Extended Data Figure 10). The association we show here of this eastern genetic character with the cultural material of the Gravettian of Buran-Kaya III, which has been compared to nearby contemporaneous Early Upper Palaeolithic assemblages from the Caucasus^11,15^ indicated in Figure 3, collectively support an eastern advance of AMHs during the EUP into Europe through the Caucasus as has been previously proposed based on archaeological evidence alone^7,8,16^. Such a population would have had to have split from the settlers of Central Europe and the Mediterranean prior to their acquisition of the Common West Eurasian component as represented by Villabruna. In this scenario, the technical adaptations required for the challenging environment of the open Eastern European plains, a dryer landscape with little natural shelter, as well as possible cultural exchanges with local populations, may have played a role in the development of the Early Gravettian industry^8^. The individuals recently characterized genetically from layer C of Dzudzuana cave in the Caucasus (data not yet available), who were found to contain ancestry (both Basal Eurasian and Common West Eurasian) that was absent ca. 9,000 years earlier in Crimea, may represent more recent immigration into Eastern Europe. A higher resolution of these movements awaits genetic analysis of more EUP and MUP sites from this region.

Numerous parallels in lithic industries, such as microblade-knapping methods, backed blades, and analogous stone blade, point and tool morphology (such as the partly backed Ahmarian el Wad-points and Gravette points of Europe) have suggested earlier Near Eastern cultures as possible precursors to Gravettian techniques^7,8^. Despite both uni-parental markers being shared between BuranKaya3A and Epipalaeolithic Natufians in the Near East, we were unable to detect extensive genome-wide allele-sharing between BuranKaya3A and the Natufians in our analysis. We note, however, that the high level of Basal Eurasian ancestry in Natufian genomic sequences (38-54%^25^) limits our sensitivity when comparing populations lacking this component. For example, negative D-statistics seen in Figure 4 involving the Mesolithic Caucasus Hunter-Gatherer (CHG) individuals Kotias and Satsurblia^37^, as well as the Natufians and the Iranian Hotu^25^, are due to the Basal Eurasian content reported for these individuals, which appears in the statistic as an affinity toward the outgroup Mbuti. Also, given the more than 20,000-year age difference between BuranKaya3A and the Natufians, it is unlikely that the Epipalaeolithic Natufians are the best surrogates for EUP Near Eastern populations. While influences from the Ahmarian can be seen in the Gravettian^17^, it is the Early Ahmarian that has separately been proposed as a source for the wave of AMHs bringing the Proto-Aurignacian west, possibly by way of the Balkans, into Central and Southern Europe beginning prior to the Campanian Ignimbrite eruption^13^. Populations associated with the Early Ahmarian and the Ahmarian may thus suggest better candidate source populations to investigate further the hypothesis of a population split behind separate routes in the settling of Europe, the Balkans/Mediterranean to the west and the Caucasus to the east, each associated with accompanying industries (Proto-Aurignacian and Early Gravettian, respectively).

The shared ancestry, as well as cultural similarities, demonstrated between the settlements at Buran-Kaya III, later Gravettians in Central Europe, and to some extent, Sunghir, suggest a broad and long-lasting network of social exchange in the EUP across Eastern and Central Europe, from the Eastern European plains to the Danubian corridor. Given this background, the appearance of the Gravettian in Central Europe in the MUP, where it later would blossom, is likely to have resulted from this input from the east. While we show that the Gravettian, *sensu lato*, was not practiced by a single genetically uniform cluster across all *facies*, the close genetic relationship between BuranKaya3A and the Kostenki individuals raise further questions as to the origin of the local culture termed Gorodtsovian, which is found at Kostenki 12 and unknown outside the Kostenki-Borshchevo region. Given the genetic affinities we report, the previous assignment by some authors of the Late Streletskian industry of Sunghir being a local *facies* of an “Eastern” Gravettian^31,32^ should lead to a closer examination of the possible influences underlying the appearance of the Gorodtsovian, with appreciation for the impact that climate, the specific landscape of the site, and site-specific activities may have had on the individual tool requirements of the assemblages^8^. Alternatively, the Gorodtsovian and other local UP industries of Eastern Europe may represent distinct cultures practiced on the Eastern European plains, and the genomic affinities between the individuals from Kostenki, Sunghir and BuranKaya3A may show only a relationship to a common source population among the occupants of Eastern Europe branching more recently than those present in Western Europe.

This study, the genomic analysis of the oldest AMH from an archeologically defined context, demonstrates an underlying genetic continuity between manufacturers of various *facies* of the Gravettian spanning ca. 9,000 years. A geographical divergence among groups entering Europe more than 37,000 years ago is supported by the finding that the earliest appearance at Buran-Kaya III is associated with a population unadmixed with the Common West Eurasian component already present in Europe to some degree, and is thus distinct. Regional features such as micro-laminar industries and Gravette points found in EUP assemblages in the Caucasus, both archeologically comparable and contemporary with Buran-Kaya III, suggest the Caucasus as a possible route for this diffusion, and a role of these industries in the development of the Early Gravettian. A more comprehensive understanding of both genomic information and archaeological assemblages of UP sites in the Caucasus and Near East will allow more precise identification of the origins of both this population and, potentially, the Gravettian.

## Methods

### Dating

All radiocarbon dates were recalibrated using the software OxCal v4.3.2 based on the IntCal13 calibration data set^38^. The calibrated dates are rounded to 5.

### Sample handling, DNA extraction, library construction, and sequencing

A human parietal fragment was excavated aseptically from layer 6-1 during the 2009 excavation season at Buran Kaya III (see Supplementary Text). All pre-amplification sample preparation was performed in the dedicated ancient DNA facility using decontamination and clean-room protocols as described in Bennett, et al^39^. All buffers and solutions were prepared using water decontaminated by gamma-irradiation (8 kGy). After first removing the surface of the areas to be sampled with a sterile scalpel, between 47 and 114 mg of bone powder was recovered from four different places of the bone using a variable-speed drill at low speed to reduce overheating (Dremel, Mount Prospect, IL, USA). Two of these samplings were each divided into two equal portions, one of which was subject to phosphate buffer pre-treatment as described in Korlevic, *et al*^40^. Phosphate buffer washes for each sample were collected and combined for DNA purification. Both phosphate and non-phosphate buffer treated samples, including reagent-only mocks, were then incubated in 1.5 mL LoBind microcentrifuge tubes (Eppendorf, Hamburg, Germany) with 1 mL 0.5 M EDTA, pH 8.0 (Sigma-Aldrich, St. Louis, MO), with 0.25mg/mL proteinase K (Sigma-Aldrich) and 0.05% UV-irradiated Tween-20 (Sigma-Aldrich), at 37°C for 24 H. Following incubation, all tubes were centrifuged at maximum speed for 10 min, and supernatant was mixed with 10 times its volume of “2M70” binding buffer (2 M guanidine hydrochloride and 70% isopropanol) in a 15 mL tube and passed through QIAquick silica columns (Qiagen, Hilden, Germany) using 25 mL tube extenders (Qiagen) and a vacuum manifold (Qiagen) as described^39,41^. 2M70 binding buffer has been shown to retain the smaller DNA fragments lost during purification with traditional binding buffers^42^. Columns were washed twice with 1 mL PE Buffer (Qiagen) then transferred to a micro-centrifuge and dried by spinning 1 minute at 16,100 × g, turning tubes 180° and repeating. DNA was eluted in a total of 60 μl of 10 mM EBT (Tris-HCl pH 8.0 containing 0.05% Tween-20) performed in two elutions of 30 μl each, by spinning 16,100 × g for 1 minute after a 5-minute incubation.

For the screening step, single-stranded libraries were constructed using either 2 μl (for screening) or 6 μl (for mitochondrial capture) of the eluted DNA, including mocks of all treatments, water only samples, and a positive control oligo following the protocol of Gansauge, *et al*.^43^ using the splinter oligonucleotide TL110, and eluting in 50 μl EBT. Either 40 μl (for mitochondrial capture) or 4 μl (for screening) of each library was used for bar-coding amplification using dual-barcoded single-stranded library adapters^44^ as primers in the following 100 μl volume reaction: 10 μl 10x PCR Buffer + MgCl2 (Roche, Basel, Switzerland) 0.4 μM of each primer, 80 μM dNTPs (Roche), 15 units of FastStart Taq (Roche). Reactions were heated 95°C for 5 min, followed by 35 cycles of 95°C 20 s, 53°C for 45 s, 68°C for 45 s, and then 68°C for 5 min. Heteroduplexes that could confound size selection were resolved by diluting the PCR product 1:5 in a 100 μl reaction containing 20 μl of the initial reaction, 8 μl PCR Buffer + MgCl2, 0.4 μM of standard Illumina primers P5 and P7, and 80 μM dNTPs, and amplified a single cycle of 95°C for 1 minute, 60°C for 2 min and 68°C for 5 min. Products were then purified and size-selected using NucleoMag beads (Macherey-Nagel, Düren, Germany) for two rounds of purification/size selection according to the supplied protocol at a ratio of bead solution 1.3 times the reaction volume and eluted in 30 µl EBT. Purified libraries were quantified using a Nanodrop ND-1000 spectrophotometer (Thermo Fisher Scientific, Waltham, Massachusetts, USA), Bioanalyzer2100 (Agilent, Santa Clara, California, USA), Qubit 2.0 Fluorometer (Thermo Fisher Scientific), and qPCR reaction. 46 to 148 ng of DNA were enriched for human mitochondrial sequence in two rounds of capture using 1200 ng of biotinylated RNA baits reverse transcribed from human mitochondrial PCR products (courtesy of L. Cardin and S. Brunel.) following the protocol described in Massilani *et al.*^45^, but with four changes: (1) for hybridization and wash steps, 60°C was used instead of 62°C. (2) DNA/RNA-bait solution was incubated 96 H instead of 48 H. (3) Elution of the bead-bound enriched DNA was performed with a 5 minute incubation in 30 µl EBT at 95°C followed by a magnetic bead separation and the transfer of the eluate to a new tube rather than a 0.1 N NaOH elution followed by silica column purification. (4) All post-capture amplifications were performed for 35 cycles followed by a heteroduplex resolution step as described above. Enriched DNA was then quantified as above, and products from all libraries were pooled in equimolar amounts and sequenced on an Illumina MiSeq using a v3 reagent kit for 2×76 cycles, substituting primer CL72 for the Read1 sequencing primer as described^44^.

Eight additional libraries along with positive and negative controls were made from 3-8 µl each of the remaining extract BK_A4B, which had the highest relative endogenous DNA content. It should be noted that this extract was derived from the portion of the cranial fragment which included the suture. Libraries were prepared as described above except one library was first treated with USER enzyme (New England Biolabs, Ipswich, Massachusetts, USA) for 30 min to remove deaminated cytosine damage. Prior to the barcoding amplification, a 6-cycle amplification of pre-barcoded libraries was performed using 45 μl of each library with 45 μl OneTaq 2X Master Mix with Standard Buffer (New England Biolabs), and 0.1 μM internal primers CL72^44^ and CL130^40^ with the above PCR conditions. Three pairs of different dual-barcoded adapters were then used to amplify 20 μl of each amplified library, followed by purification and size selection as described above, which allowed the later pooling of two enrichment protocols and non-enriched DNA from the same library. An average of 1.2 μg of DNA to 1 μg RNA-baits was used for each mitochondrial enrichment, as above, however, an additional alternative “touch-down” hybridization protocol consisting of 60°C for 12 H, 59°C for 12 H, 58°C for 12 H, 57°C for 12 H, and 56°C for 48 H was tested for each library. For select libraries, an alternative wash protocol described in Fu *et al*.^46^ was also tested. Neither of these alternative protocols had a substantial impact on the results obtained. Enriched and shotgun libraries were then pooled separately and size selected on an E-Gel SizeSelect 2% agarose gel (Thermo Fisher). Enriched libraries were then sequenced on an Illumina MiSeq, as above, and shotgun libraries were sequenced on an Illumina NextSeq using a NextSeq 500/550 High Output Kit v2 (2×75 cycles).

### Data analysis

Paired-end sequencing results were merged and adapters trimmed using leeHom^47^, and reads were then aligned to the human genome (hg37d5) with BWA (v0.7.12) aln, parameters −n 0.01 −l 0, followed by samse^48^. Reads shorter than 28 bp long were then removed directly from sam files with an awk command^39^. PCR duplicates were removed using MarkDuplicates (v2.9.0)^49^ and reads mapping with quality less than 25 were removed with SAMtools (v1.7)^50^. Reads mapping to the nuclear genome with a mapping quality score of 25 or greater were locally realigned around known indels using GATK (v3.7-0) IndelRealigner^51^. Following this step, mapped reads less than 35 bp containing indels were removed^52^. This step reduced aberrant SNP calls due to spurious alignments of short fragments in our dataset, where the base did not match either of two expected alleles, from 1.1% to 0.6%. In comparison, raising the minimum length of all reads to 30 bp reduced aberrant SNP calls to 0.4%, but reduced informative SNPs by 18.4%, demonstrating the utility of retaining a short minimum read length while excluding short indel-containing reads for this sample. All libraries showed extremely poor preservation of genetic material. The average fragment length of mapped reads after the above treatment was 38 bp, and the first position C>T transition rate from damage was 55% at the 5’ end of the molecule as calculated by mapDamage (v2.0.6)^53^ (Extended Data Figure 12).

Reads mapping to the mitochondrion were used to determine the posterior probability for contaminating modern human sequences with Schmutzi^54^ and found to have a distribution maximum at 1%. A mitochondrial consensus sequence was called from the majority of bases at each position using Geneious (v8.1.9)^55^, which exactly matched that generated from Schmutzi. The 5’-most 100 bp of this consensus sequence was duplicated at the 3’ end and used as a reference for a new alignment using the enriched reads. The haplogroup was called using Phy-Mer with Build 16 rCRS-based haplogroup motifs^56^ and verified by manual analysis of sequence changes. A Bayesian tree of 25 ancient and 3 modern mitochondrial sequences, excluding the hypervariable regions, was constructed using MrBayes^57^ using a GTR+i+G nucleotide substitution model, which gave the lowest log-likelihood for the tree out of all models tested (GTR and HKY with all combinations of 4 invariant sites and gamma distributions), agreeing with the results of JModelTest2^58^. The chain was run for 1,100,000 iterations, subsampled every 200 after discarding a 9% burn-in period and visualized using FigTree v. 1.4.3 (http://tree.bio.ed.ac.uk/software/figtree). A list of sources for the mitochondrial sequences is given in Table S5.

Genetic sex was determined using the ratio of sequences aligning to the X and Y chromosomes, given as the R_y_ value, as described in Skoglund, *et al.*^26^. An additional calculation of the ploidy of the X chromosome using read counts mapping to the X chromosome to the autosomes, (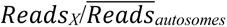) was also performed. SNPs informative for the Y-chromosome haplogroup were identified using Yleaf (v1.0)^59^ for reads with mapping quality scores of both 10 and 20 (Supplementary Data 1).

### Neanderthal content

Neanderthal ancestry was calculated using the ancestry informative SNP method^1^ calculated separately for Neanderthal- or AMH-derived SNPs on four libraries prepared from two independent extractions, treated either with or without UDG. The combined libraries overlapped a total of 6,252 SNPs reported to have derived in either the Neanderthal (2,538 SNPs) or AMH (3,714 SNPs) lineages^60^. The combined libraries averaged 3.5% (SD 0.0074) Neanderthal ancestry. To test the sensitivity of these SNPs to determine Neanderthal ancestry, this SNP subset was then used to recalculate Neanderthal ancestry from several previously reported UP genomes^1^. The results agreed with previously reported values (Table S4).

### SNP calling and f-statistics

BuranKaya3A bam files for all barcoded libraries were merged with SAMtools merge^50^ and PCR duplicates removed with MarkDuplicates^49^. SNPs were called for positions overlapping with the combined SNP panels from Fu *et al*.^1^ using SAMtools mpileup^50^, requiring a SNP base quality score of 30 or greater, and choosing one allele at random when necessary with pileupcaller^61^. This resulted in 27,740 SNPs out of 2,990,848 being called for BuranKaya3A. To monitor the impact cytosine deamination may have on the SNP calls and resulting analyses of ancient samples, alternative datasets removing all C to T or G to A transitions were generated and all statistical analyses were performed on both data sets. Results with >30% disagreement between the two datasets were excluded from the analyses, but both datasets are included in the supplements.

To calculate f-statistics, data from 52 previously reported genomes (in either fastq or bam format) were downloaded and realigned to the human genome (hg37d5), and SNPs were called following the identical pipeline used for BuranKaya3A, with the exception that diploid SNP calls were retained for high-coverage individuals used as an outgroup in *f*3 analyses (list of samples and references given in Table S6). Modern humans used in statistical analyses are from Mallick, et al.^62^. “Mbuti” is a population of three modern Mbuti individuals. *f*3-statistics and D-statistics were computed using ADMIXTOOLS^29^ qp3Pop (v412) qpDstat (v712), respectively. Standard error was estimated using a block jackknife with 0.050 centiMorgan blocks. Full analyses performed and results for *f*3(*x*, *y*; Mbuti) and *D*(*w*, *y*; BuranKaya3, Mbuti) are given in Table S7 and Supplementary Data 2, respectively.

### Data Availability

Sequence data generated in this study will be made available upon publication.

## Acknowledgments

We thank Olivier Gorgé for assistance with DNA sequencing. We thank also the National Academy of Sciences of Ukraine for permission to excavate at Buran-Kaya III, and all the team members of the 2001 and 2009–2011 excavation seasons. EAB was supported by the CNRS and the Labex “Who am I”. SPr was supported by the Foundation Fyssen. SPe was supported by the French National Research Agency (ANR-05-JCJC-0240-01). The field work was supported by the Muséum National d’Histoire Naturelle (MNHN, Paris), the CNRS and French Ministry of Foreign Affairs. The paleogenomic facility obtained support from the University Paris Diderot within the program “Actions de recherches structurantes”. The sequencing facility of the Institut Jacques Monod, Paris, was supported by grants from the University Paris Diderot, the Fondation pour la Recherche Médicale (DGE20111123014), and the Région Ile-de-France (11015901).

## Author contributions

EMG, SPr, and SP initiated the project, EMG and EAB designed the study. EAB performed the laboratory work, EAB, TG, and EMG analysed the data, LC performed aseptic excavation of the sample. SPr performed the anthropological analysis. AY, SP and SPu provided archaeological data. EAB wrote the manuscript with input from EMG, TG, SPr, SP, LC, and AY.

---

**Extended Data Figure 1.**
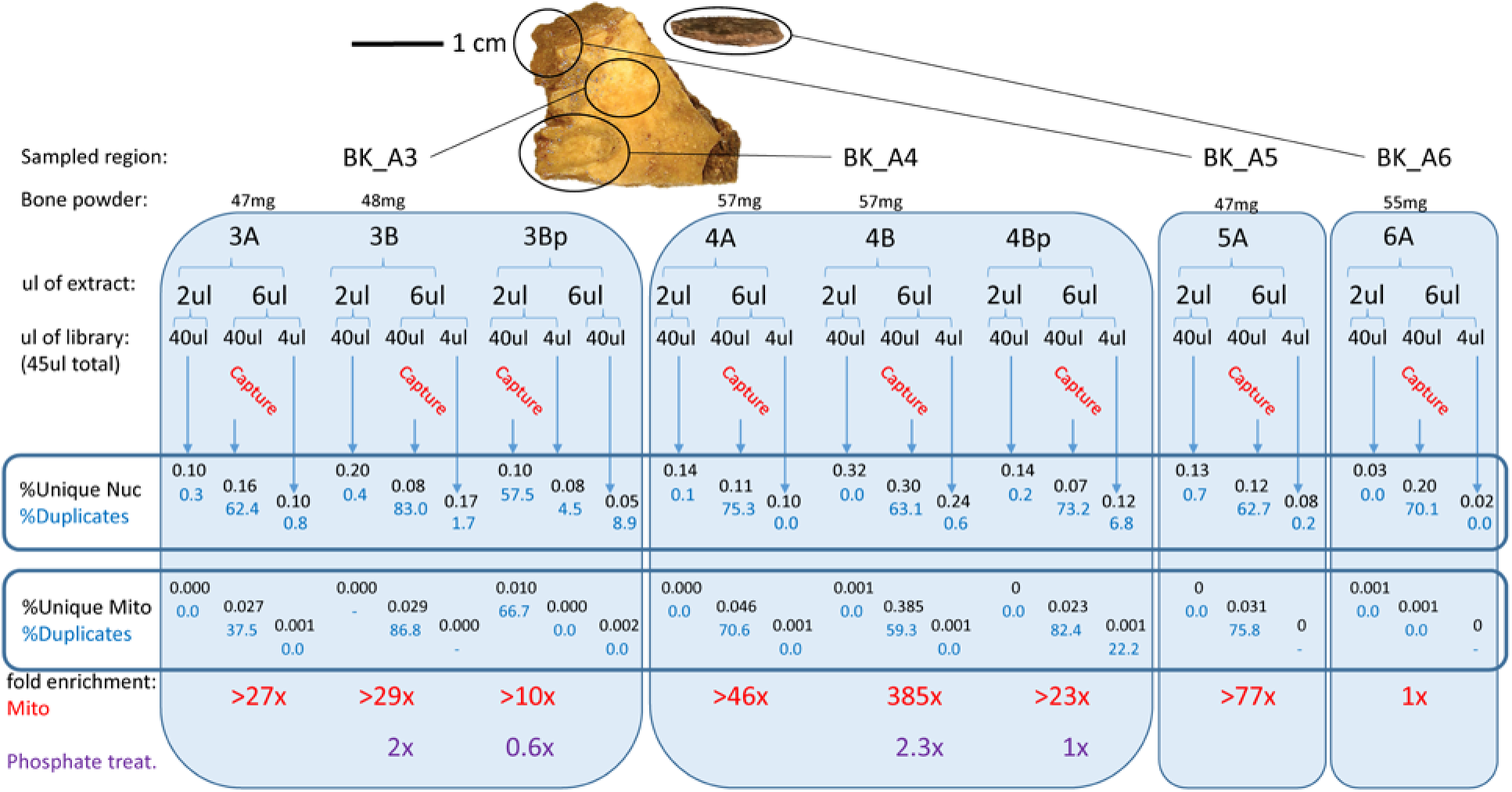
Flowchart of screening methods and results of mitochondrial capture and phosphate enrichment for each library. Results show the percent of unique reads out of all reads 28 nucleotides or greater mapping to the nuclear and mitochondrial references as well as the percent of PCR duplicates for each library. The suffix of sample numbers is as follows, A = no phosphate treatment, B = phosphate treated, Bp = phosphate treatment buffer. Extract BKA4B was selected for subsequent analysis. Original bone sample and regions used for each library is shown above. It has been noted that the region containing the darker suture yielded the best results.

**Extended Data Figure 2.**
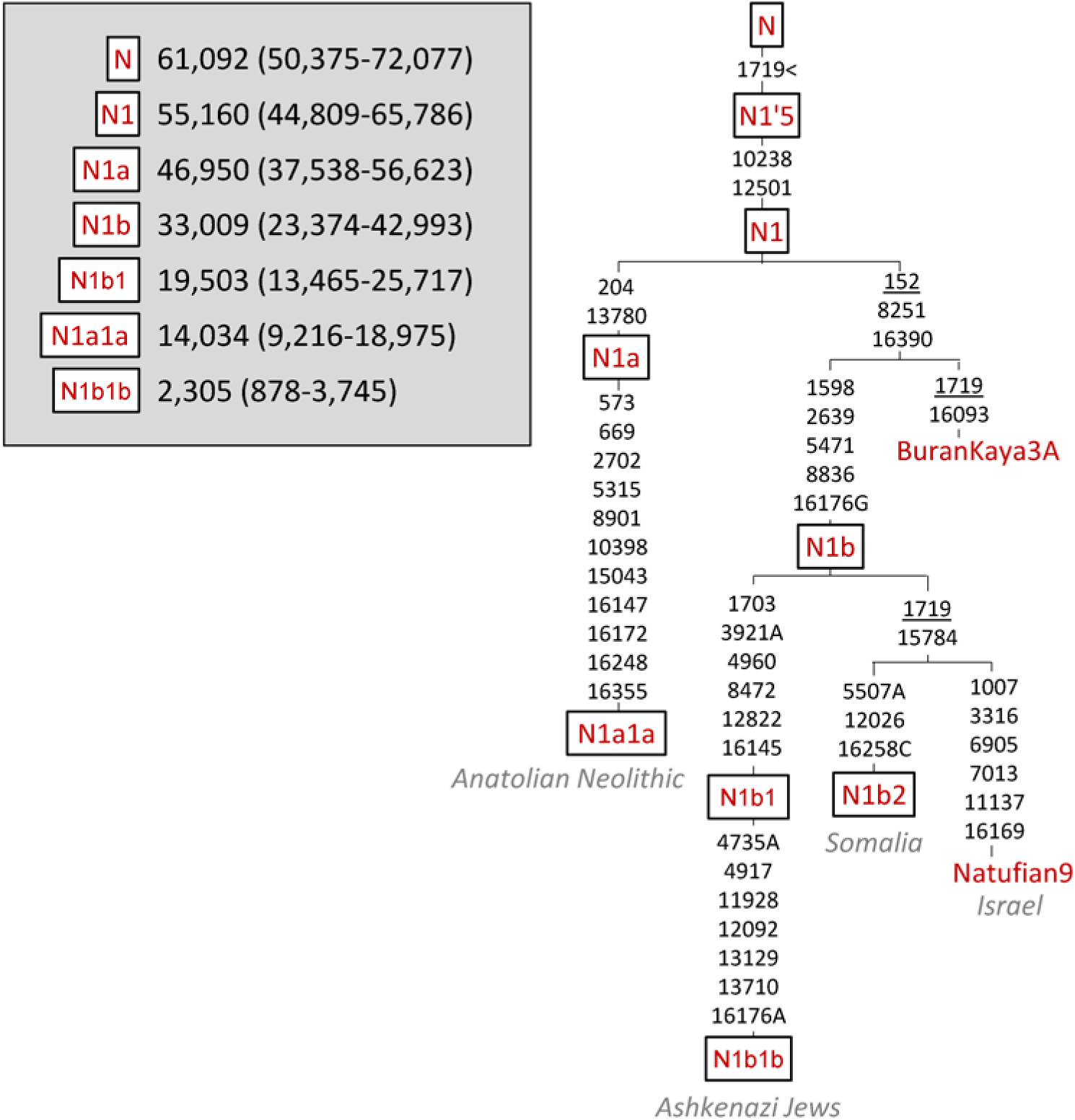
Detailed mutation map of mitochondrial N clade with the positions of BuranKaya3A, the Epipalaeolithic Natufian (Natufian9), and lineages discussed in the text. Grey italics represent the population or geographical region where the clade is prominent. Inset indicates the maximum likelihood estimation of TMRCA values of the nodes with 95% CI from Fernandes et al.^22^. Mutation tree calculated by mtphyl version 5.003 (https://sites.google.com/site/mtphyl/home) compared to the rCRS. Underlined positions represent back mutations.

**Extended Data Figure 3.**
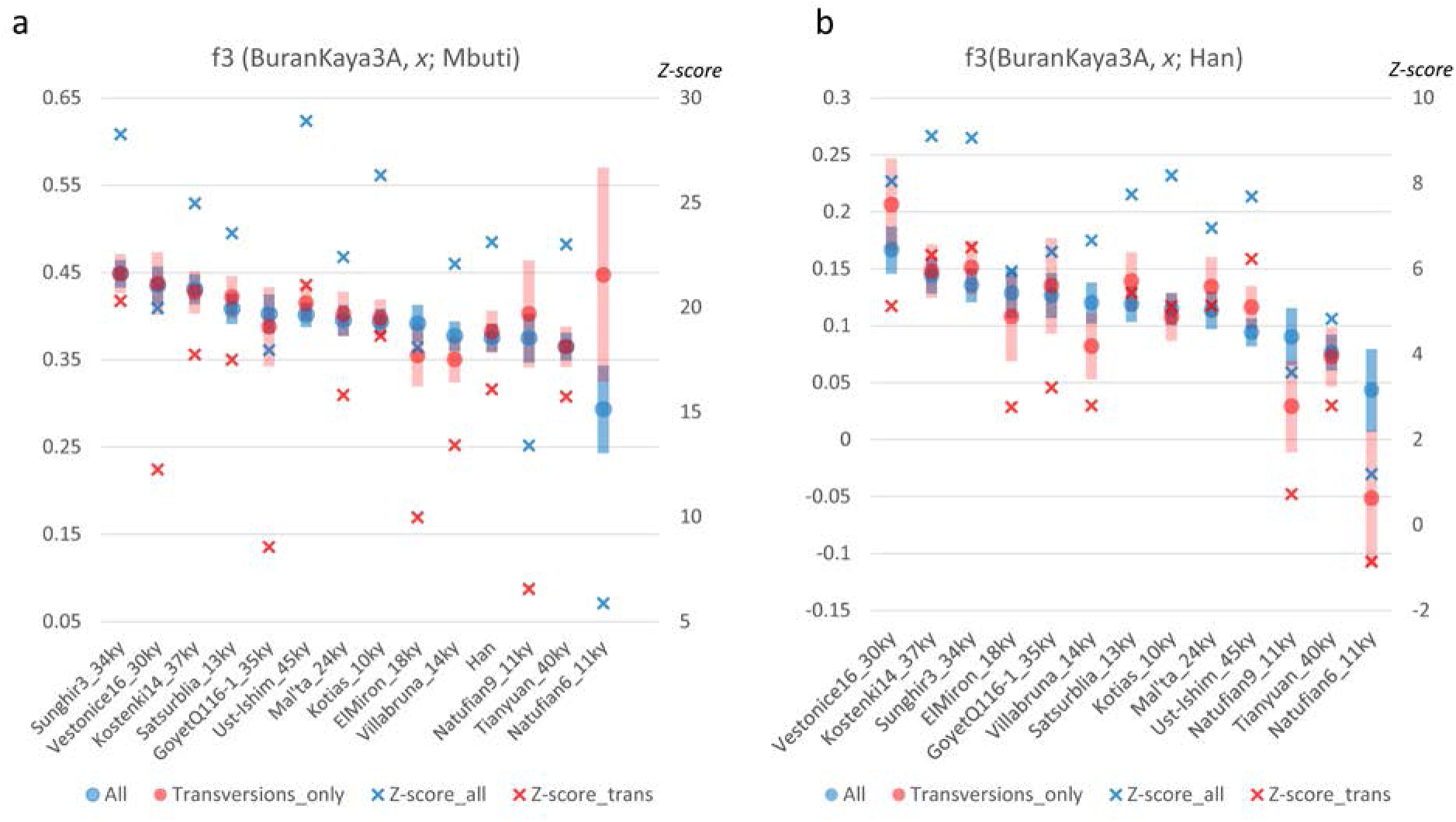
Outgroup *f*3-statistics results showing the degree of shared alleles between BuranKaya3A and high-coverage UP Eurasians for outgroups Mbuti (a), Han (b). Left axis: *f*3-statistic corresponding to circles with error bars. Right axis: Z-score corresponding to “x”. Results for all SNPs are in blue, transversions only in red. The degree of overlap between blue and red shows the degree of agreement between the two datasets for a given sample combination. Approximate ages are appended to the names. Boxed names indicate individuals associated with a Gravettian context. Error bars = one standard error.

**Extended Data Figure 4.**
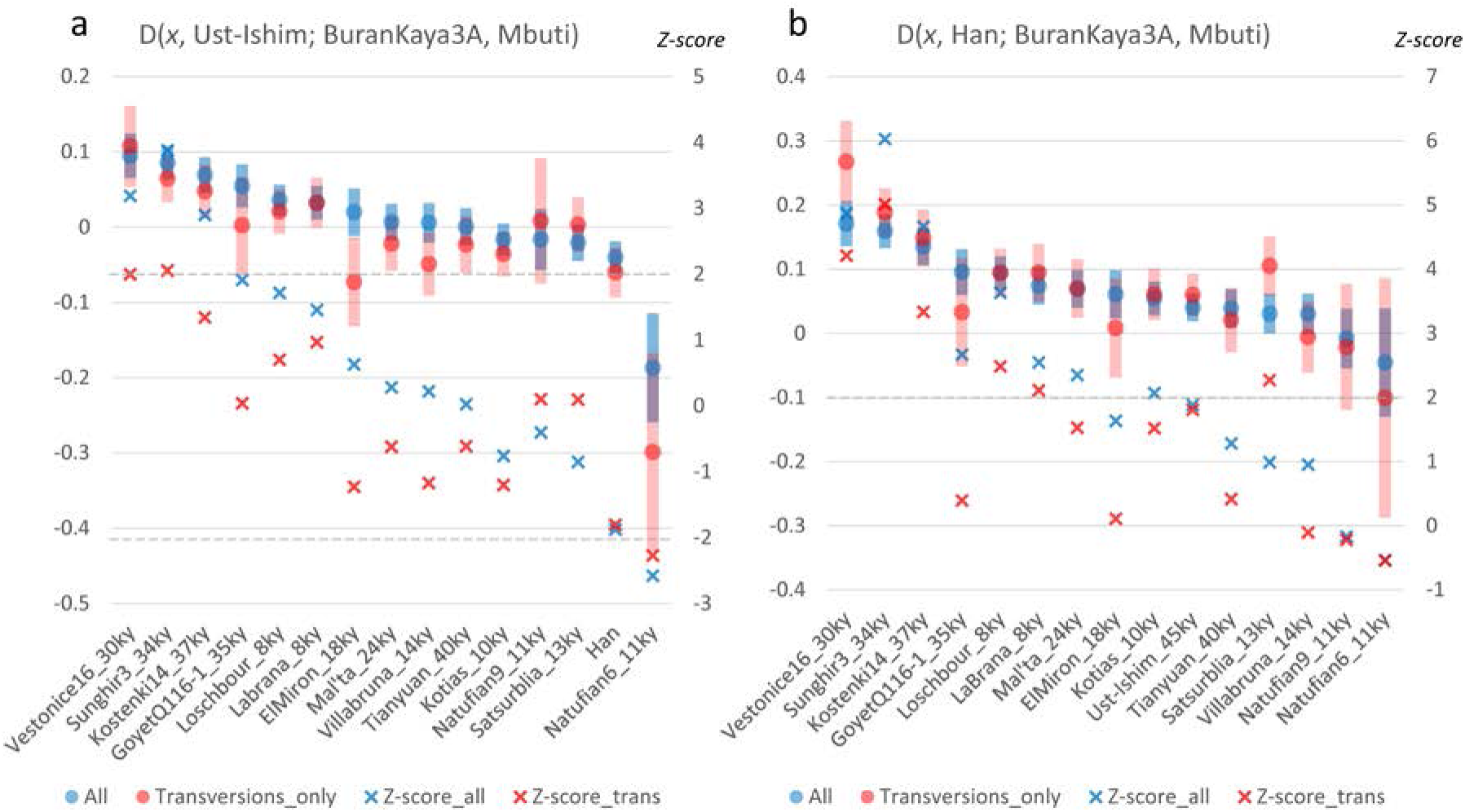
D-statistics results for a) *D*(*x*, Ust-Ishim; Burankaya3A, Mbuti) and b) *D*(*x*, Han; BuranKaya3A, Mbuti) where *x* = high coverage UP individuals. Left axis: D-statistic corresponding to circles with error bars. Right axis: Z-score corresponding to “x”. Results for all SNPs are in blue (shown also in Figure 4), transversions only in red. The degree of overlap between blue and red shows the degree of agreement between the two datasets for a given sample combination. Approximate ages are appended to the names. Boxed names indicate individuals associated with a Gravettian context. The values for Kotias, Satsurblia, and the Natufians are distorted in (a) due to Basal Eurasian content in these samples. Error bars = one standard error.

**Extended Data Figure 5.**
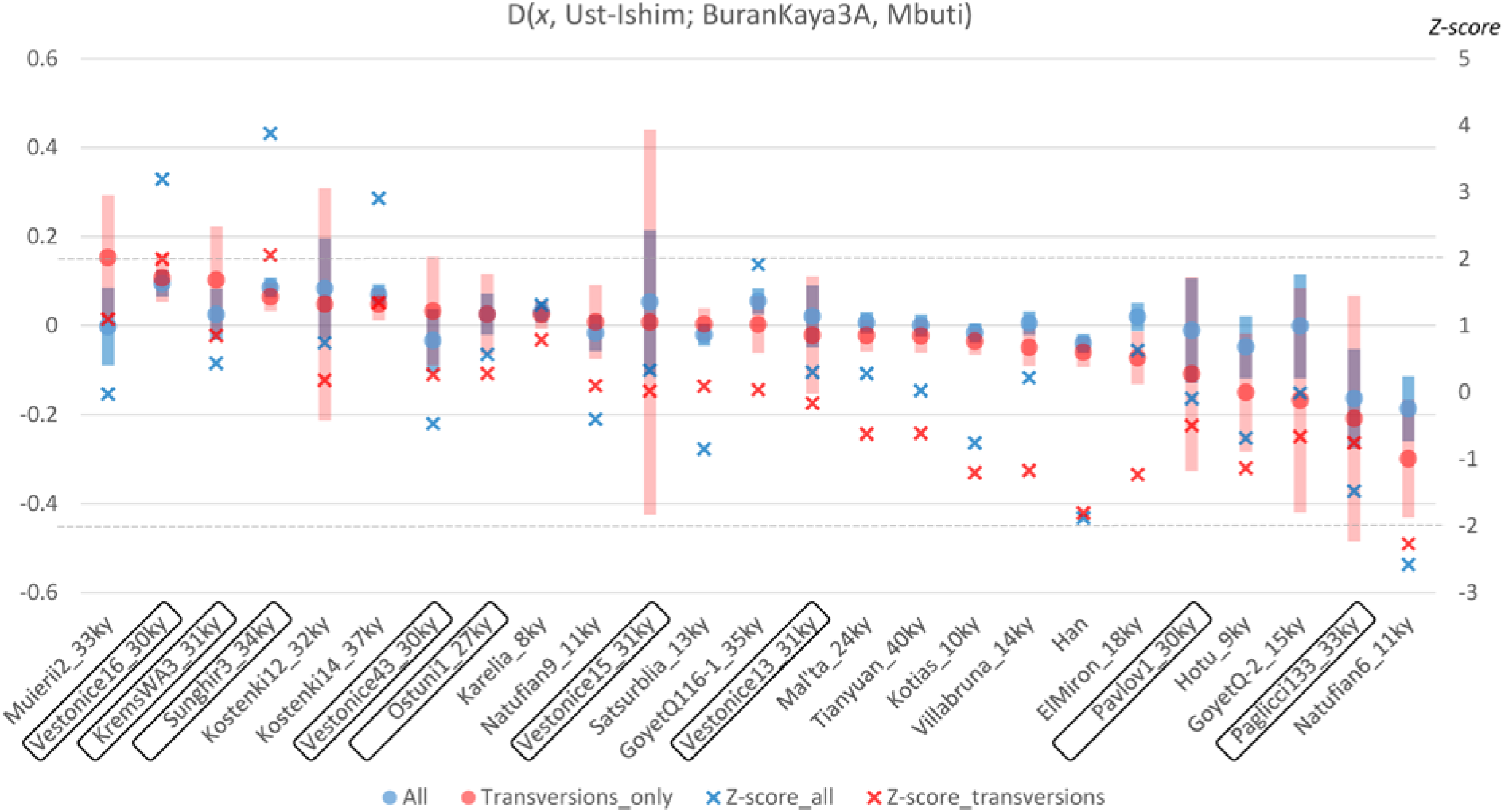
D-statistics results for *D*(*x*, Ust-Ishim; Burankaya3A, Mbuti) where *x* = both high and low coverage UP individuals and a modern East Asian (Han). Only samples using at least 100 transversion-only SNPs for the calculation are shown. Left axis: D-statistic corresponding to circles with error bars. Right axis: Z-score corresponding to “x”. Results for all SNPs are in blue, transversions only in red. The degree of overlap between blue and red shows the degree of agreement between the two datasets for a given sample combination. Approximate ages are appended to the names. Boxed names indicate individuals associated with a Gravettian context. The values for Kotias, Satsurblia, and the Natufians are distorted due to Basal Eurasian content in these samples. Error bars = one standard error.

**Extended Data Figure 6.**
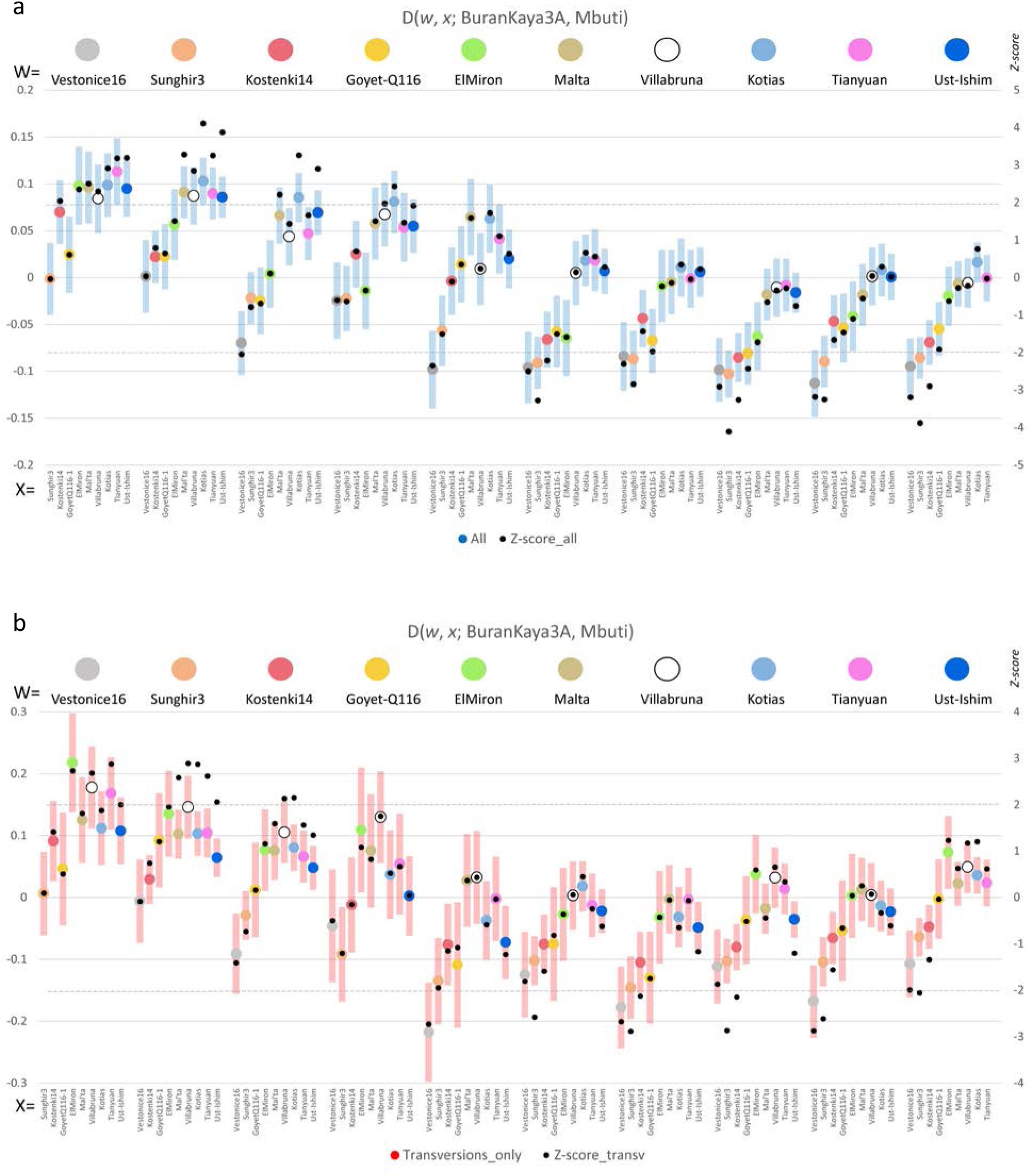
D-statistics results for *D*(*w*, *x*; BuranKaya3A, Mbuti) where *w* and *x* are high-coverage UP Eurasians. More positive values show an excess of allele sharing of *w* over *x* with BuranKaya3A. Left axis: D-statistic corresponding to circles with error bars. Right axis: Z-score corresponding to black dots. Significance (Z-score > 2) is indicated by horizontal dashed lines. The values for Kotias are distorted due to Basal Eurasian content in this sample. Error bars = one standard error. a) Results for all SNPs, b) results for transversions only.

**Extended Data Figure 7.**
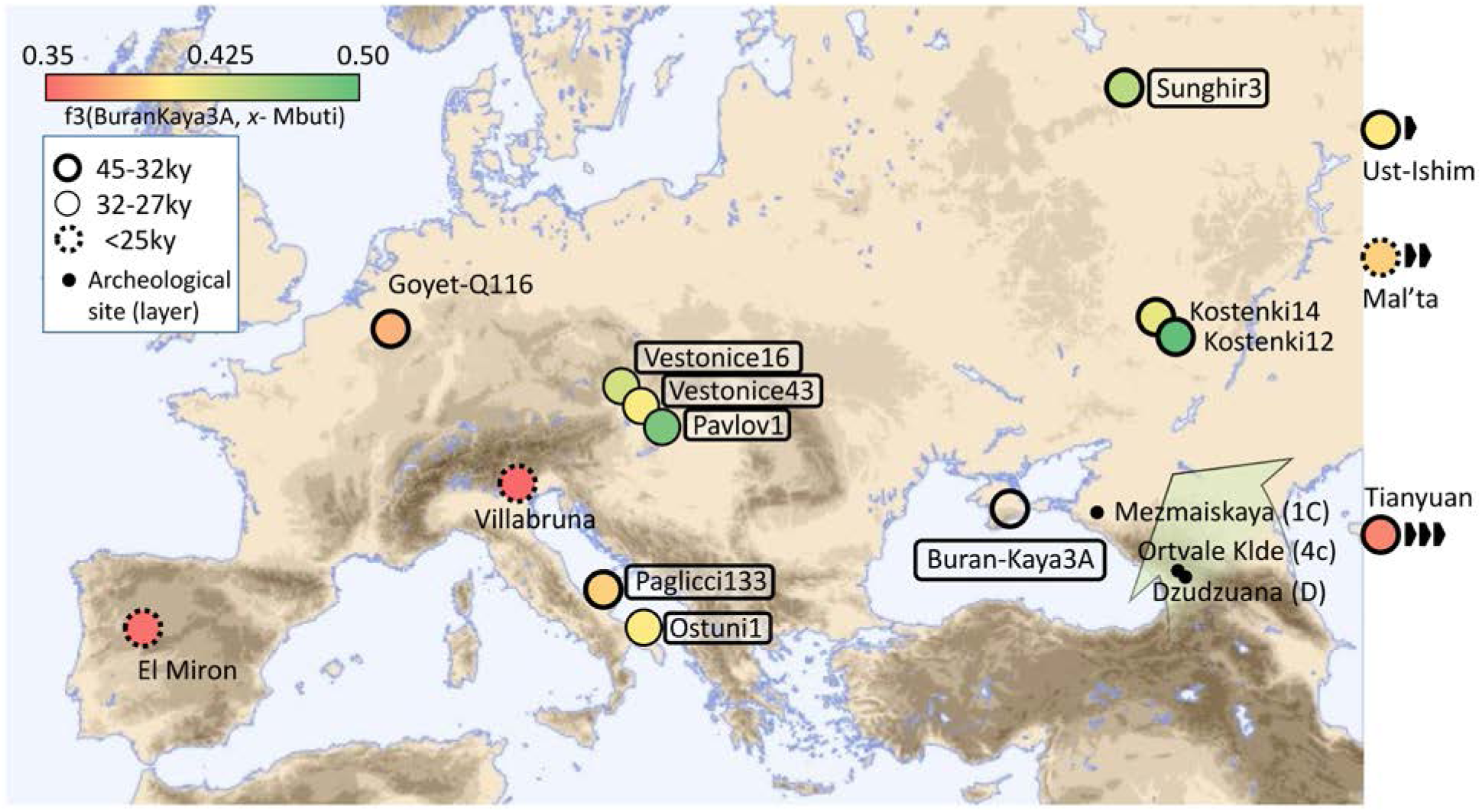
Location and heatmap of *f*3(BuranKaya3A, *x*; Mbuti) with archaeological sites mentioned in the text. Values, as in Figure 3, but adding results from *x* including the lower-coverage samples whose calculations made use of at least 50 transversion-only SNPs and were within 30% agreement of values calculated using all SNPs. Boxed text indicates samples associated with a Gravettian archaeological context (including Sunghir3, alternatively described as Streletskian). Non-European samples are given along the right margin at their approximate latitudes, their relative distances indicated by arrows. Archaeological sites for which no human genetic data are available that contain micro-laminar industries comparable with Buran-Kaya III at contemporaneous layers (in parentheses) indicated by black dots. A broad green arrow shows the proposed EUP route introducing the Early Gravettian into Europe, suggested by the similar features of these assemblages and the lack of a Common Western Eurasian genomic component in BuranKaya3A.

**Extended Data Figure 8.**
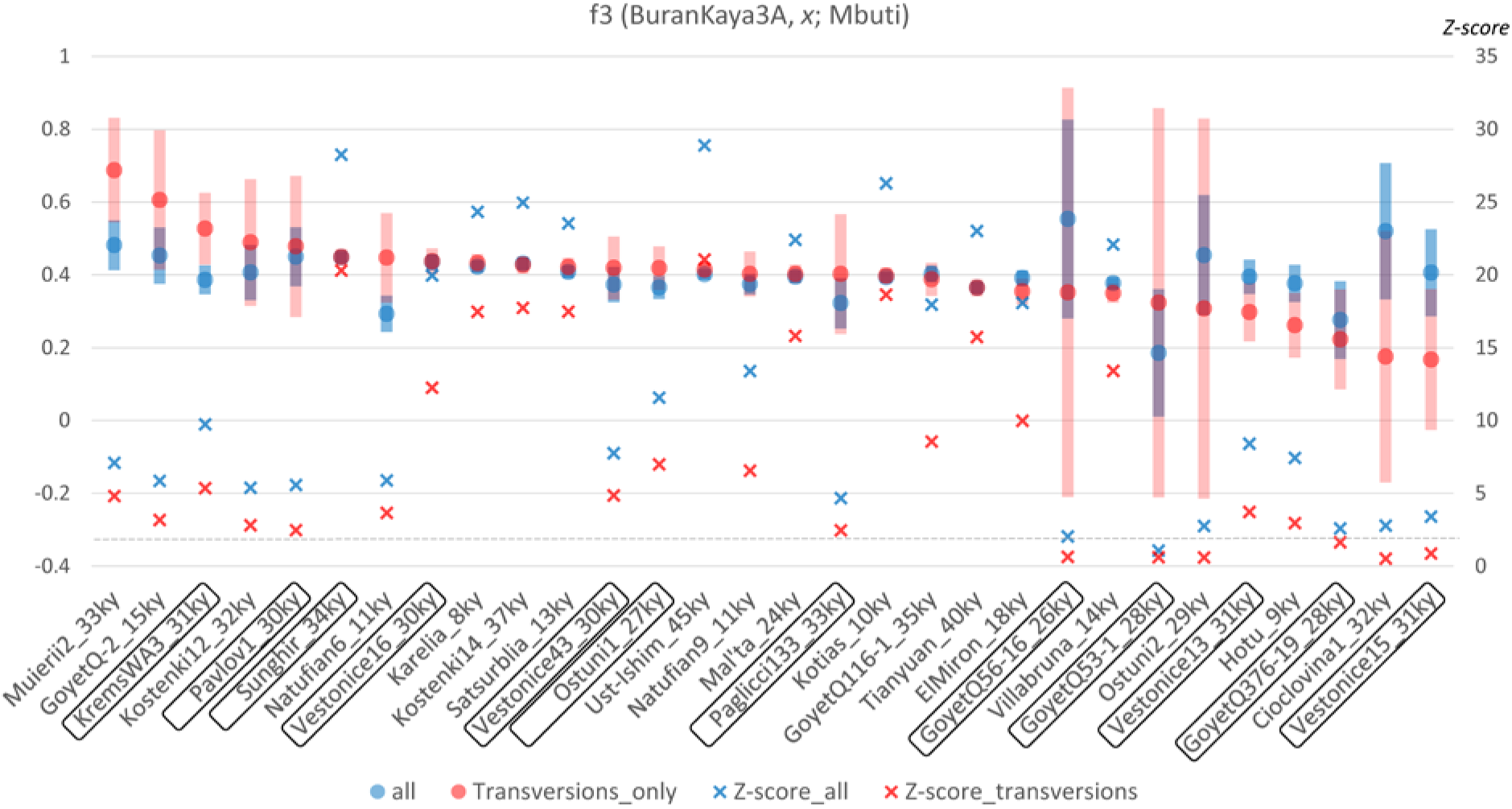
Outgroup *f*3-statistics results showing the degree of shared alleles between BuranKaya3A and UP Eurasians for outgroups Mbuti, *f*3(BuranKaya3A, *x*; Mbuti), as in Extended Data Figure 3A but including all samples for which values could be calculated. Left axis: *f*3*-statistic* corresponding to circles with error bars. Right axis: Z-score corresponding to “x”. Results for all SNPs are in blue, transversions only in red. The degree of overlap between blue and red shows the degree of agreement between the two datasets for a given sample combination. Approximate ages are appended to the names. Boxed names indicate individuals associated with a Gravettian context. Error bars = one standard error.

**Extended Data Figure 9.**
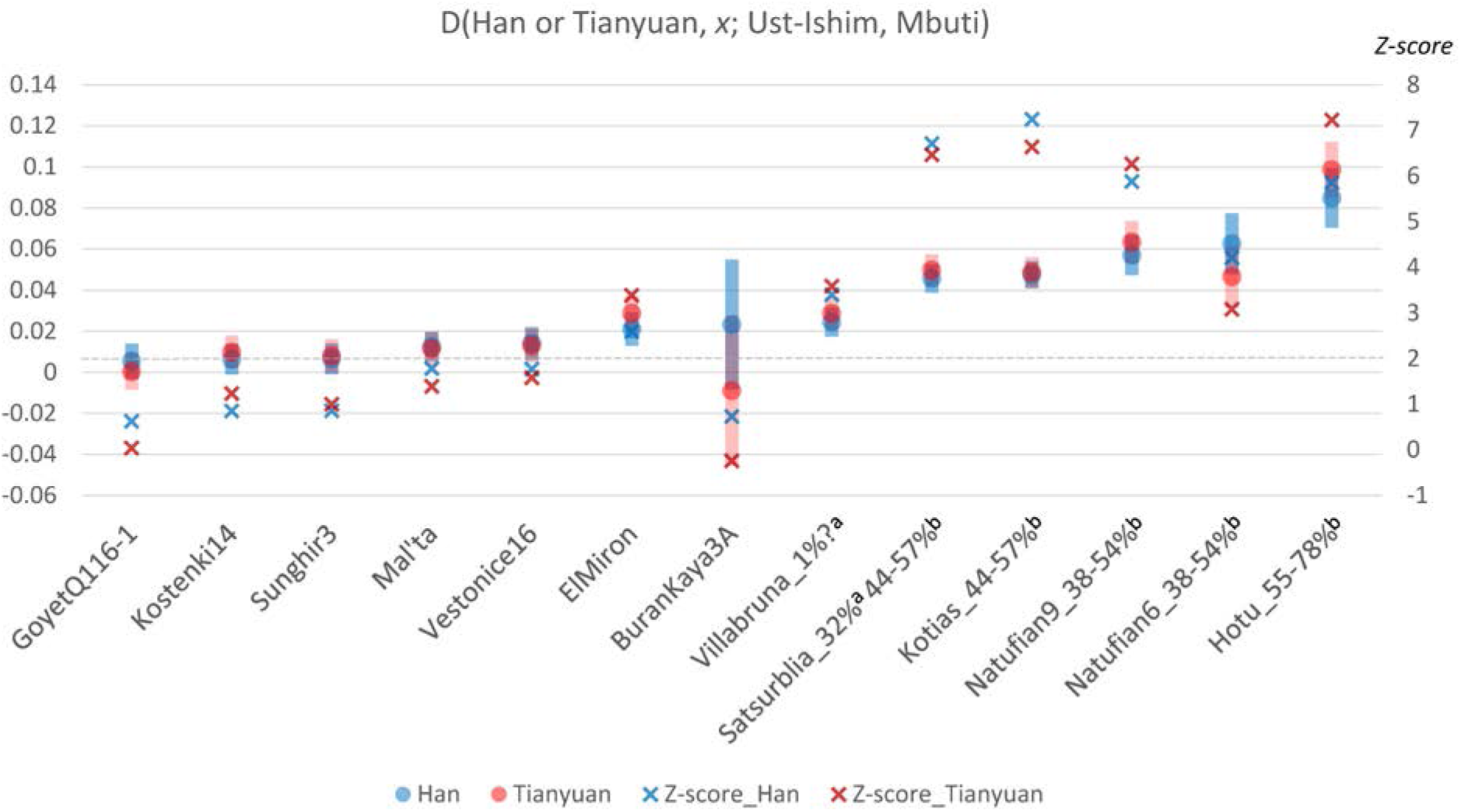
Relative Basal Eurasian content as demonstrated by the statistic *D*(*Modern or Ancient East Asian*, *x*; Ust-Ishim, Mbuti)^1^ using the transversions only SNP dataset. Left axis: D-statistic corresponding to circles with error bars. Right axis: Z-score corresponding to “x”. *x* =Han, blue, *x*=Tianyuan, red. Significance (Z-score = 2) is indicated by a horizontal dashed line. Estimated Basal Eurasian content percentages given as suffixes to the names (a) estimated from ADMIXTUREGRAPH^1^, (b) one standard error range estimated from the f4 ratio^25^. Error bars = one standard error.

**Extended Data Figure 10.**
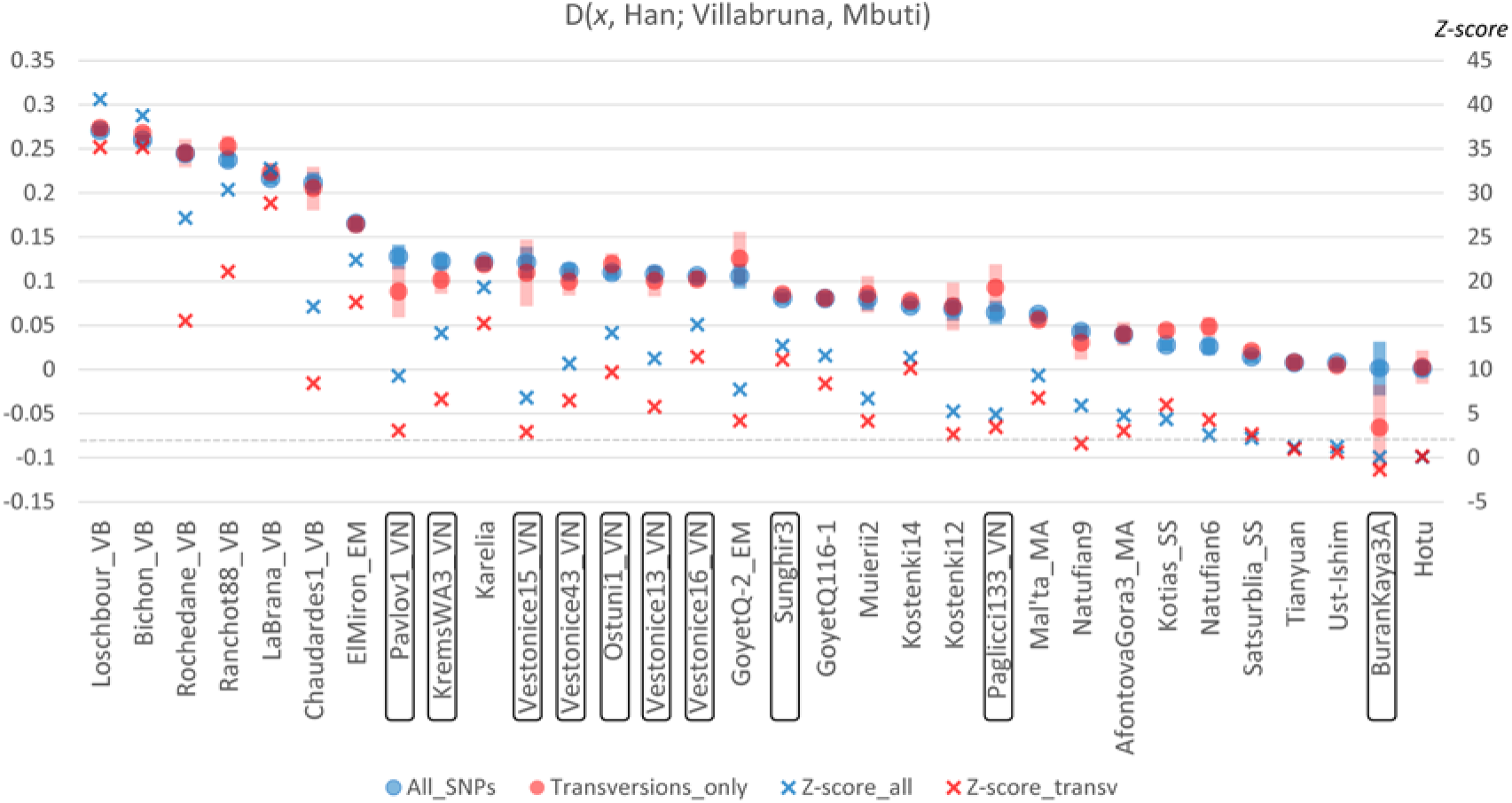
Relative Common West Eurasian content (as represented by Villabruna) given as *D*(*x*, Han; Villabruna, Mbuti) for ancient samples (*x*) using more than 1,000 SNPs for the calculation. More positive values signify an excess of shared alleles between *x* and Villabruna as compared to Han. Reported genetic clusters^1^ given as suffixes to the names: VN, Vestonice; EM, ElMiron; VB, Villabruna; SS, Satsurblia; MA, Mal’ta. Boxed names indicate individuals associated with Gravettian contexts. Left axis: D-statistic corresponding to circles with error bars. Right axis: Z-score corresponding to “x”. Results for all SNPs are in blue, transversions only in red. The degree of overlap between blue and red shows the degree of agreement between the two datasets for a given sample combination. This value will be distorted for samples having Basal Eurasian content (see Extended Data Figure 9). Significance (Z-score = 2) is indicated by a horizontal dashed line. Error bars = one standard error.

**Extended Data Figure 11.**
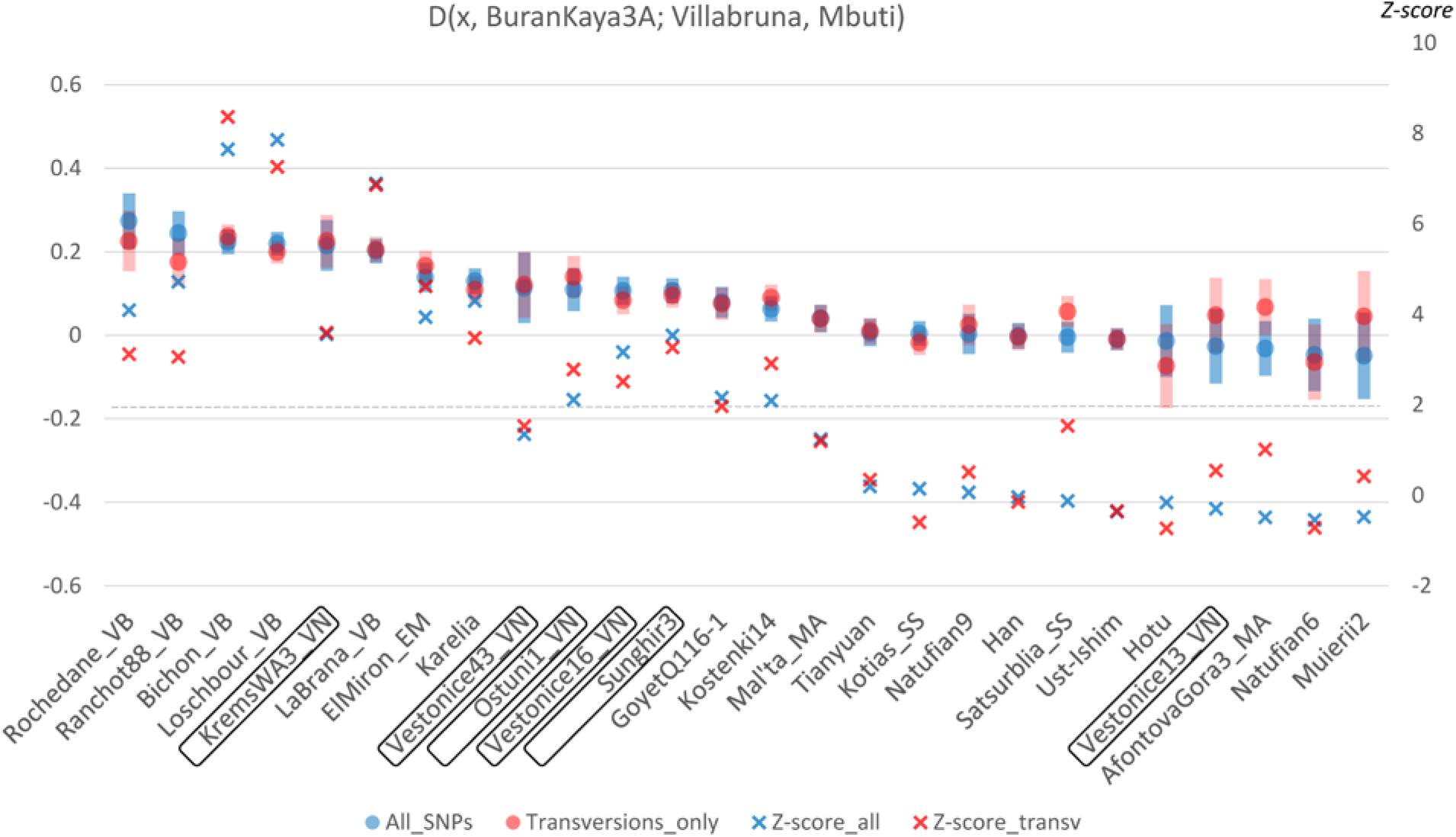
Measuring the excess of shared alleles between Common West Eurasian (as represented by Villabruna) and *x* over BuranKaya3A using the D-statistic *D*(*x*, BuranKaya3A; Villabruna, Mbuti). More positive values signify an excess of shared alleles between *x* and Villabruna compared to BuranKaya3A. Reported genetic clusters^1^ given as suffixes to the names: VN, Vestonice; EM, ElMiron; VB, Villabruna; SS, Satsurblia; MA, Mal’ta. Boxed names indicate individuals associated with Gravettian contexts. Left axis: D-statistic corresponding to circles with error bars. Right axis: Z-score corresponding to “x”. Results for all SNPs are in blue, transversions only in red. The degree of overlap between blue and red shows the degree of agreement between the two datasets for a given sample combination. This value will be distorted for samples having Basal Eurasian content (Extended Data Figure 9). Significance (Z-score = 2) is indicated by a horizontal dashed line. Error bars = one standard error.

**Extended Data Figure 12.**
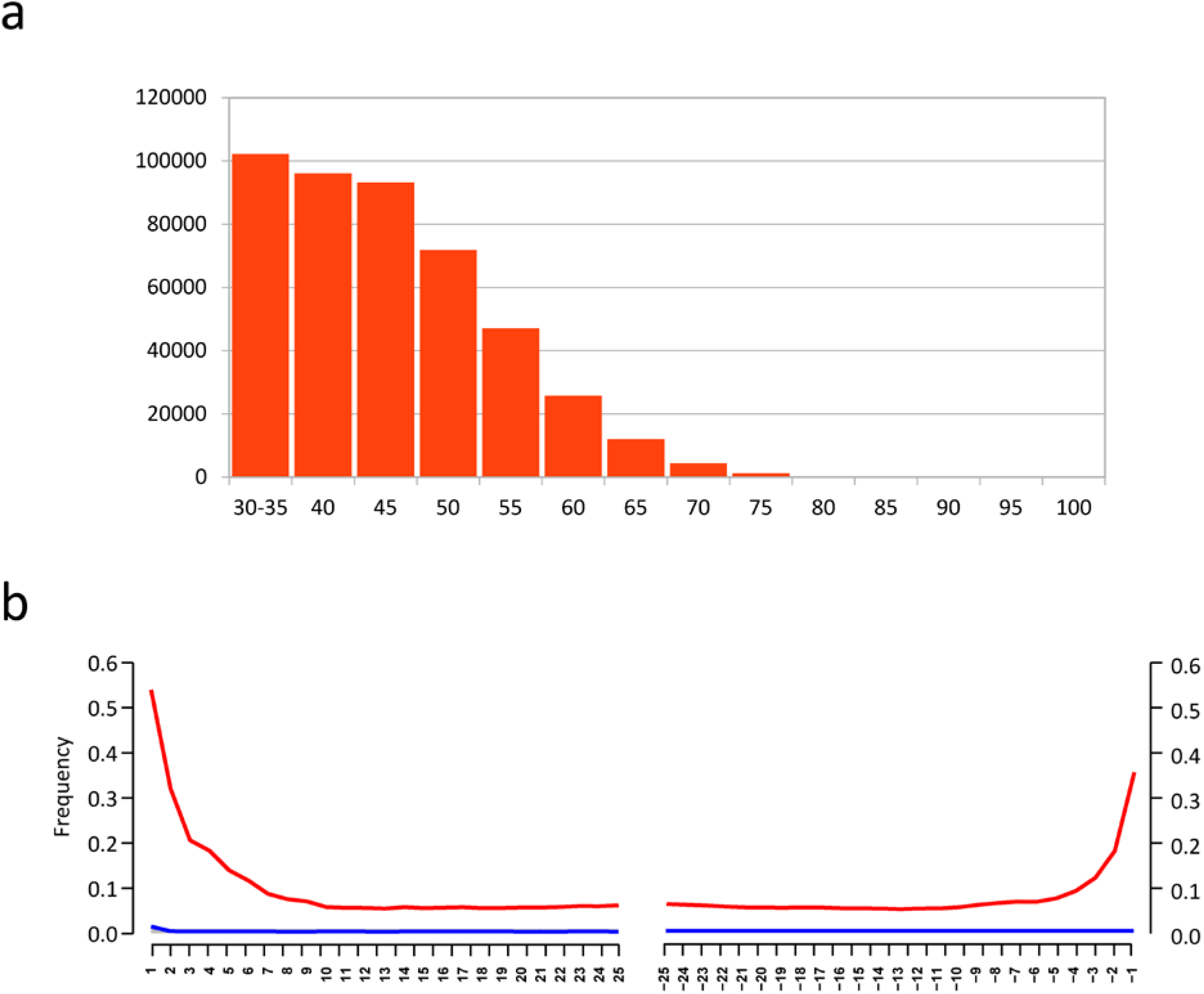
a) Size distribution of recovered BuranKaya3A DNA fragments mapping to the nuclear genome. b) The frequency of C to T mismatches due to cytosine deamination of the first 25 nucleotides from the 5’ and 3’ ends of the DNA molecules. Graph generated by mapDamage (v2.0.6)^53^.

