## Supplementary Text and Supplementary Tables S1-S7 for "The origin of the Gravettians: genomic evidence from a 36,000-year-old Eastern European"

### Table of Contents

**Supplementary Text**

1. Cultural context of the Gravettian technocomplex 2

2. Buran Kaya III site, chronological, cultural and anthropological information 5

3. Strategy of collection and sampling protocol 12

Supplementary Text References 15

**Supplementary Tables**

Table S1. Comparative $^{14}C$ dates 21

Table S2. Shotgun library results 22

Table S3. Y SNP summary with Palaeolithic comparisons 23

Table S4. Neanderthal content comparison 24

Table S5. Mitochondrial sequence references 25

Table S6. Genome sequence references 27

Table S7. *f3*(BuranKaya3A, *x*, Mbuti) results 29

Supplementary Tables References 31

**Supplementary Text 1. Cultural context of the Gravettian technocomplex**

**By Stephane Péan**

UMR 7194 (HNHP), MNHN/CNRS/UPVD, Muséum national d'Histoire naturelle, Alliance Sorbonne Université, Institut de Paléontologie Humaine, Paris, France

### Definition and general chronology

The Gravettian is defined as a mid-Upper Palaeolithic, European-wide, technocomplex of cultural traditions[1–3], named after the site La Gravette (France), where specific lithic straight-backed points (*pointe de la Gravette*) were first described[4].

The Gravettian lithic industry can be characterized by elaborated, light, regular and precise *débitage* and straight-backed pointed blade tools, which probably represent projectile implements[2,5]. The Gravettian cultural facies present also characteristic osseous industry artefacts, personal ornaments, portable art (e.g. female figurines), or spatially organized archaeological structures made from stones or bones interpreted as dwelling structures at some sites.

The Gravettian cultures in Central Europe have been divided into eight regional *facies*[2] and three main chronological phases[6–9]: early phase, characterised by predominant leaf-shaped points and *fléchettes,* in the Middle Danube valley (Austria, Moravia in Eastern Czech Republic); middle phase, characterised by abundant Gravette points and microgravettes and the appearance of new microlithic and para-geometric implements, in Lower Austria, Moravia, Slovakia and northeastern Hungary, culturally related to Eastern European sites in the Middle Dniester and Don valleys; and late phase, characterised by shouldered points, in the middle Danube valley (Lower Austria, Moravia, western Slovakia), Southern Poland, middle Dniester valley, Upper Dnieper valley and the Middle Don and Seym valleys, in eastern Europe.

The Gravettian technocomplex chronologically spans from 32 to 21 ka $^{14}C$ BP(ca. 36-23 ka cal BP)[10,11] (see below for early Gravettian sites). During late MIS 2, Gravettian evolves towards Epigravettian *facies*, which last until 13.5 ka $^{14}C$ BP (16.4-16.1 ka cal BP).

**Origin of the Gravettian cultural *facies***

Central Europe includes Gravettian settlements dated as early as ca. 32-30 ka $^{14}C$ BP: in South Poland, at 32,400 ± 650 $^{14}C$ BP (i.e.38.3-35.1 ka cal BP) and 31,000 ± 550 $^{14}C$ BP (36.1-34.0 ka cal BP) in Obłazowa cave layer VIII[12], at 31,550 ± 350 $^{14}C$ BP (36.1-34.7 ka cal BP) in Henryków 15 site layer 9[13]; in Austria, at 30,500 + 900/-800 $^{14}C$ BP (36.7-33 ka cal BP) in Willendorf II layer 5[14]; and in Ukraine, at 29,650 ± 1,320 $^{14}C$ BP (37.1-31.1 ka cal BP) in Molodova V layer IX[15].

In terms of origin, different hypotheses have been proposed. Considering an internal evolution within Europe, Kozlowski (2015) argues a polycentric rather than a monocentric origin to be more conceivable, i.e., in the middle Danube basin (as a prominent center), as well as the Dniester and Prut basins, and Crimea[16]. In this hypothesis, the Gravettian could have evolved locally from regional post-Mousterian transition industries with leaf-shaped points in Eastern Europe[17], such as the Szeletian into the Pavlovian in the Middle Danube basin[18], or similar local transitional industries into the Molodovian in the Dniester valley[19]. The Gravettian could have also evolved from the Aurignacian in the Upper Danube valley[20], the Proto-Aurignacian in the Balkans, the Uluzzian in the Mediterranean areas or the Zwierzyniecian in the areas north and east of the Carpathians[19]. In contrast, one hypothesis of an initial origin of the Gravettian in Central Europe postulates that it may have resulted from acculturation between migrating Anatomically Modern Humans importing Aurignacian cultures and local Neanderthals producing leaf point industries[5].

An external origin of the Gravettian has also been proposed linking it to cultural *facies* of the Near East and Northeastern Africa, such as the Ahmarian, Lagaman and Dabban[21].

The industry from Buran-Kaya III layers 6-2, 6-1 and 5-2, attributed to the Gravettian technocomplex[22,23], has been considered, together with the material from layers 6-5 to 6-3, as having a number of characteristics similar to Early Upper Palaeolithic assemblages from the northwestern Caucasus (Mezmaiskaya Cave, levels 1C, 1B and 1A) and southern Caucasus (Dzudzuana cave , layer D; Ortvale Klde rockshelter, layers 4d and 4c)[24].

The last $^{14}C$ analyses of the three layers 6-2, 6-1 and 5-2 from Buran-Kaya III, radiometrically dated between 32.5+250-230 and 29.4+190-180 ka $^{14}C$ BP (37.1-33.2 ka cal BP)[25,26], confirm the hypothesis of a very early occurrence of Gravettian settlements prior to 30 ka BP in Eastern Europe[16,27].

**Supplementary Text 2. Buran Kaya III site, chronological, cultural and anthropological information**

**By Sandrine Prat**

UMR 7194 (HNHP), MNHN/CNRS/UPVD, Alliance Sorbonne Université, Musée de l'Homme, Paris, France

**Chronological, cultural context and settlement patterns**

Buran-Kaya III is a rock shelter located on the eastern part of Crimea, in the Belogorsk region, in the middle basin of the Burulcha River (4 km south from the city of Aromatne). This site was discovered in 1990 by A. Yanevich and excavated initially until 2001 by a team directed by A. Yanevich (Neolithic to Palaeolithic layers) and A. Marks (Middle and Early Upper Palaeolithic layers) with the participation of V. Chabai, Y. Demidenko, K. Monigal, M. Otte and Y. Yamada, followed by new fieldwork conducted by A. Yanevich and S. Péan between 2009 and 2011. This rockshelter unearthed an exceptional stratigraphic sequence from the Middle Palaeolithic to the Neolithic which lead to multiple studies[22,23,25,26,28–47].

Buran-Kaya III is a key site for understanding the Middle to Upper Palaeolithic transition and the arrival and dispersal of Anatomical Modern Humans (AMHs) in Europe as well their potential biological and/or cultural interactions with Neanderthals. During the excavation season of 2001 and 2009-2011, this site has yielded a Middle Palaeolithic layer (Micoquian, Kiik-Koba type, layer B) located above an Early Upper Paleolithic layer (Streletskian or eastern Szeletian, layer C) and below three Aurignacian layers (layers 6-5 to 6-3), followed by three Gravettian layers (layers 6-2, 6-1 and 5-2) and one Swiderian layer (layer 4), all densely layered across 1 to 2 meters of depth.

The stratigraphical and radiochronological analyses[26] had emphasized an early chronological framework for the Middle to Upper Paleolithic period in southeastern Europe. These studies show the presence of Early Upper Paleolithic (early Streletskian or eastern Szeletian, layer C) populations in Crimea, before 40.0 ka cal BP confirming a previous dating of 44.3–38.5 ka cal BP (36.7 ± 1500 ka BP, OxA-6868) of the Streletskaya or eastern Szeletian layer[30] and providing early evidence of this transitional industry. Based on the new radiocarbon dates, the late Middle Palaeolithic settlements (Micoquian layer, layer B) range between 35,390 +290/-270 and 37,700 ± 900 BP[26] (43,545 and 39,300 cal BP), showing that no Neanderthal settlements occurred at Buran-Kaya III after the Campanian Ignimbrite eruption around 39 ka cal BP. The two Upper Paleolithic cultural traditions, related to the Aurignacian and Gravettian complexes (layers 6-5 to 5-2), were successively present in Crimea during a relatively short time span between 39.4 and 34.1 ka cal BP. During the following period of about 20,000 years, no depositional processes occurred or were preserved and the site was presumed abandoned. The Final Paleolithic populations (Swiderian, layer 4) settled in Crimea between 11.8 and 11.3 ka cal BP, i.e. during MIS 1 (Fig. S1).

The layers (6-2, 6-1 and 5-2) yielded a large collection of human remains, and more than 28,000 lithic remains were discovered during the 2001 field season, representing a large spectrum of tool types (e.g. burins, end-scrapers, and backed microliths). The lithic industries exhibited a high percentage of microliths (more than 80%) and some microgravettes[23], which are consistent with the Gravettian technocomplex. They do not present Dufour or pseudo-Dufour blades, often associated with Aurignacian technocomplex, which are present in layer 6-3, 6-4 and 6-5, stratigraphically located below[23]. As shown by the sedimentological and stratigraphic analyses conducted by S. Puaud during the 2009 and 2010 field seasons, there is continuity in the stratigraphic sequence from the lower layers (Aurignacian technocomplex: 6-5, 6-4, 6-3) to the upper layers (Gravettian technocomplex: 6-2, 6-1 and 5-2). The attribution of the upper layers to the Gravettian tradition is free from material sedimentary overlap.

Concerning the bone industry, more than 60 bone tools have been discovered in the Gravettian layers corresponding to projectile points, awls, arrowheads, and assegai points[23]. The bone industry was probably imported as indicated by the absence of manufacturing waste. This evidence, combined with the zooarchaeological results[44], suggests a pattern of recurrent short-term occupations, such as seasonal hunting or temporary butchery camps, associated with exploitation of small or medium-sized mammals, such as saiga antelope (the main game exploited in these layers). The settlement of AMHs associated with the Gravettian layers of Buran-Kaya III occurred under interstadial climatic conditions, which became cooler and drier leading to an open steppe environment. The open and steppe-like periglacial environment, with a local transition zone between steppes and mountains located on the migration path of several species, provided a diversified faunal spectrum (e.g., saiga antelope, hare, red and polar foxes, reindeer, woolly rhinoceros, marmot, wild cat and brown bear)[26,44,47]. The micromammal and palynological analyses[36,48] reveal a steppe environment with a climatic evolution towards an increasing aridity as recorded in the mammalian sequence.

**Human remains**

At Buran-Kaya III, more than 160 human remains were discovered in three well-documented Gravettian layers (6-2, 6-1 and 5-2). Two of them unearthed from layer 6-1 and 6-2 have been directly dated to 31,900+240/-220 BP (36.3-35.2 ka cal BP), layer 6-1 (GrA-37938)[25], and 32,450

+250/-230 BP (37.1-35.7 ka cal BP), layer 6-2 (GrA-50457)[26]. They are among the oldest direct evidence of AMHs in Europe in a well-documented archaeological context (Gravettian *sensu lato*) [25,26]. The Gravettian specimens from Buran-Kaya III represent, together with Kostenki, Russia (Kostenki 14: 33,250 +/-500 BP; 38.7-36.3 ka cal BP[49] and Peştera cu Oase, Romania (Oase 1: 34,290 +970/-870 BP: 36.47-41.07 ka cal BP, GrA-22810)[50], the earliest occurrence of AMHs in Eastern Europe. AMHs occurred later in Western Europe, e.g., at Goyet Cave in Belgium (Goyet Q116-1: 30,880 +170-160 BP; 35.2-34.4 ka cal BP, GrA-46175)[51], in Russia, e.g., at Sunghir (Sunghir SI: 28,890 +/- 430 BP; 33.9-31.8 ka cal BP, OxA-X-2464-12; Sunghir SII: 30,100+/-550; 35.3-33.2 ka cal BP, OxA-A-2395-6; Sunghir SIII: 30,000 +/-550 BP; 35.2-33.3 ka cal BP, OxA-X-2395-7; Sunghir SIV: 29820 +/- 280 BP; 34.5-33.5, OxA-X-2462-52)[49,52] and in the Czech republic e.g., at Dolní Věstonice (Dolní Věstonice 16, associated with Gravettian tradition: 25,740 +/-210 BP (on charcoal); 30.6-29.4 ka cal BP, GrN-15277)[49].

Layer 6-1 has yielded the richest assemblage of human specimens with 150 highly fragmented remains (mostly cranial parts and teeth (95%) and several hand phalanges), corresponding to at least five individuals[45]. Due to the fragmentation of the remains, the taxonomic assignment as AMH could only be performed on an occipital bone fragment and the permanent teeth. Based on the combinations of morphological features on these anatomical elements (Figure 2) (namely, absence of an occipital “bun” or a bilaterally transverse torus; lack of a well-developed metaconid and a transverse crest on the lower premolars; lack of shovelling, labial convexity and the presence of well-developed lingual tubercles on the upper first incisors; the lack of a well-developed mid-trigonid crest and a large anterior fovea on the lower molars), these individuals from layer 6-1 are allocated to AMHs[25,47].

Isotopic analyses of bone collagen revealed that the diet of these individuals consisted mainly of terrestrial resources, mammoth meat playing a preponderant role in the protein intake, and of plant consumption[46].

Among the human remains from the layer 6-1 (2001, 2009 and 2010 field seasons), only a few bones exhibit human modifications, such as cut marks. The human skulls were intentionally selected in association with post-mortem treatments of the corpses (ritual cannibalism or a specific mortuary practice, such as post-mortem disarticulation processing of corpses for secondary disposal)[25,44,47]. Based on these taphonomic observations, the specimens from layer 6-1 represent the oldest Upper Palaeolithic AMHs from Eastern Europe showing post-mortem treatment of the corpses.

194

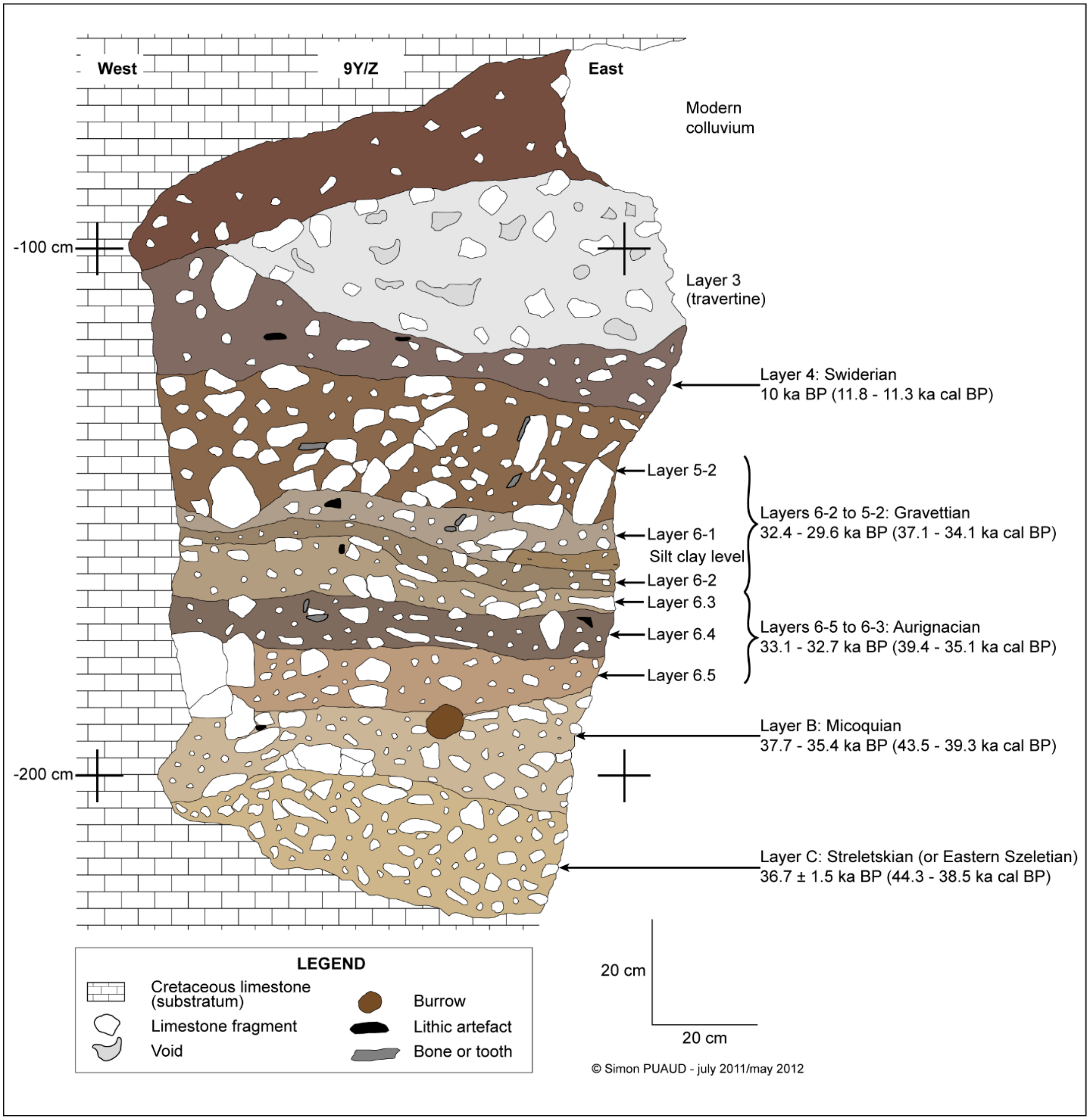


195 **Fig. S1.** Stratigraphy and chronological frameworks of Buran Kaya III site (graphic S. Puaud),

196 modified from Péan *et al.*, 2013[26].

197

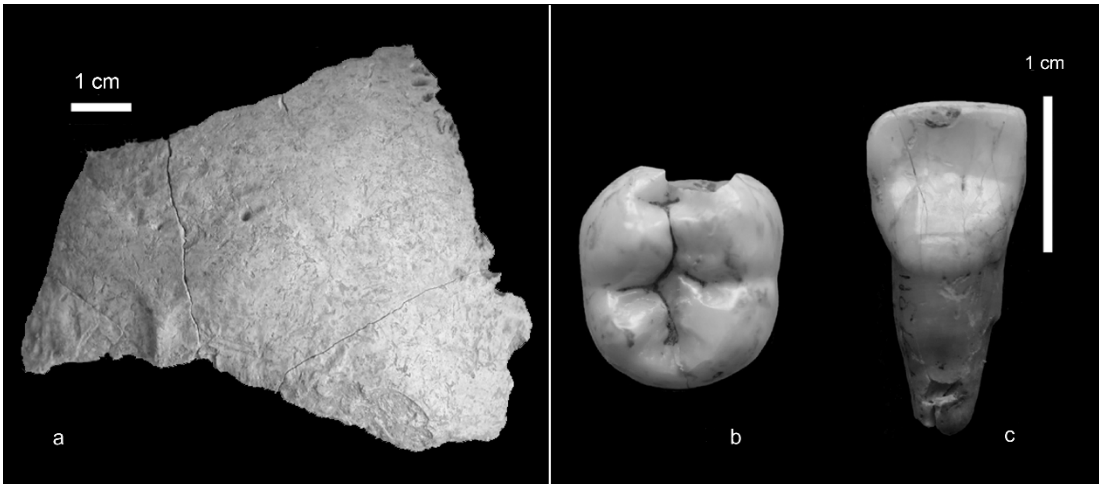


**Fig. S2.** Remains from Buran-Kaya III (layer 6-1). From left to right, a) occipital bone; b) right lower second molar; c) right first upper incisor. Scale bar = 1 cm; photos © L.Crépin/S. Prat.

### 3. Strategy of collection and sampling protocol

**By Laurent Crépin**

UMR 7194 (HNHP), MNHN/CNRS/UPVD, Muséum national d’Histoire naturelle, Alliance Sorbonne Université, Institut de Paléontologie Humaine, Paris, France

The excavations performed between 2009 and 2011 incorporated a multi-sampling strategy to better contextualize the site of Buran-Kaya III. Sampling for sedimentological, biogeochemical, and radiometric studies were performed on all Gravettian layers, and in 2009, the sampling for the palaeogenomic analyses was carried out focused on the two layers that contained most of the human remains (6-1 and 6-2). To this end, a one half-square meter section was excavated incorporating precautions against contamination: a maximum of two excavators dug the square wearing masks and gloves, and when potential human remains were discovered, one excavator put on a new full body protection suit, including masks and gloves, to perform the removal of the material using new tools (Fig. S3a).

This protocol was maintained over eight days, during which time layers 6-1 and 6-2 were entirely cleared. In total, six bones were identified and inventoried using the described procedure, of which four were certified as human remains after verification: three from layer 6-1 (two cranial fragments and one tooth) and one for layer 6-2 (a cranial fragment). These remains, separated from other archaeological remains, were transferred directly after the mission to the high containment laboratory at the “Institut Jacques Monod” (IJM), where they were analyzed and stored at -20°C. Additionally, all those working on the site during the excavation campaign provided a cheek-swab sample to control for possible contamination of the samples.

The anthropological and taphonomic study of the human remains was then carried out directly in the high containment palaeogenomic laboratory at the IJM using typical precaution measures for ancient DNA research (Fig. S3b). Finally, we decided to carry out the paleogenomics analysis on the bone fragment that we could identified anatomically in an unambiguous manner that appeared to be the best preserved with a minimum of visible alteration: artefact number N.135, square 9Z, layer 6-1.a also called here *BuranKaya3A* (Fig. S3c). This piece is a posterior parietal bone fragment presumably belonging to a mature individual, measuring a maximum of 21 x 19 mm.

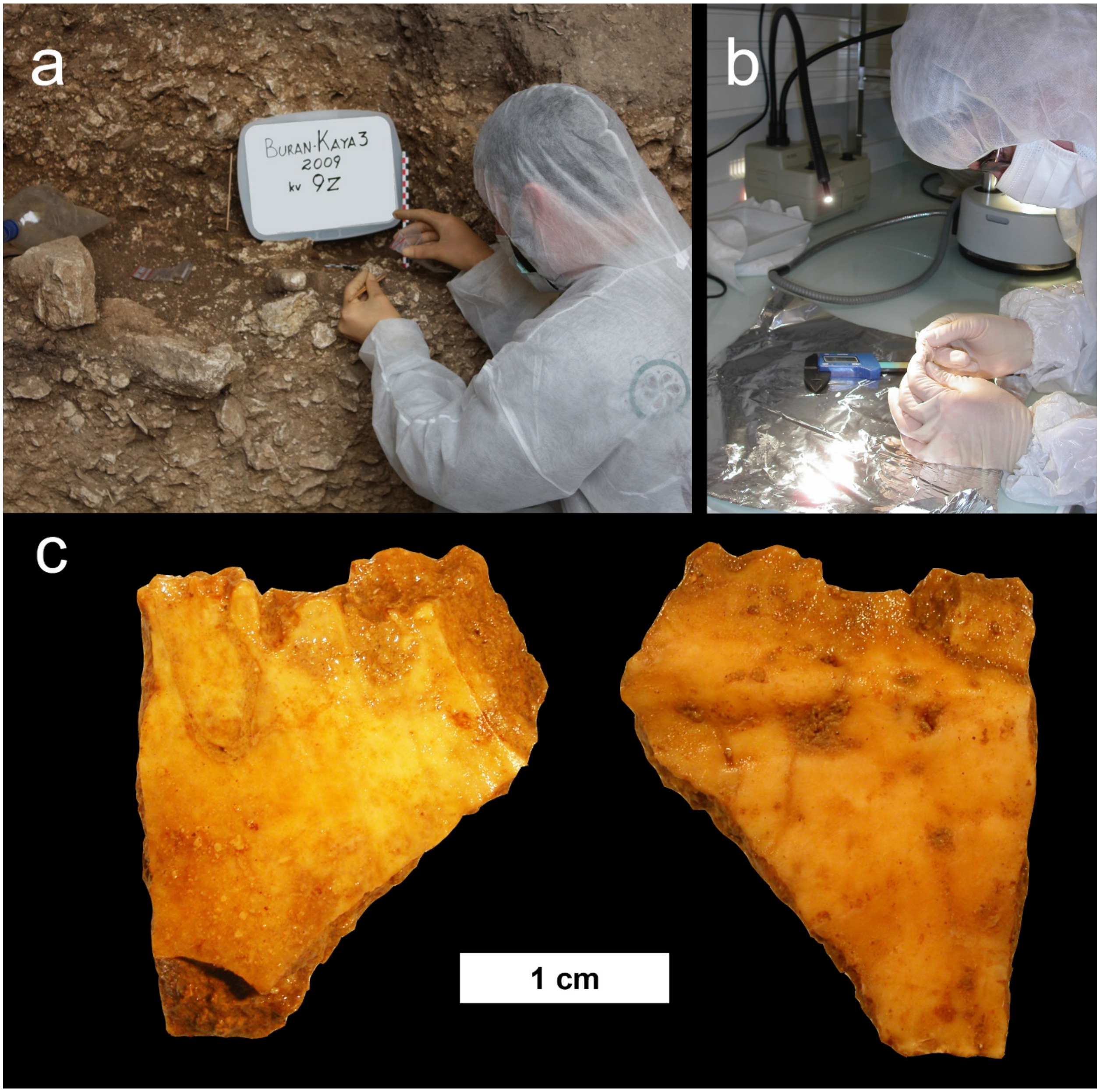


**Fig. S3.** a) Sampling of human remains during the 2009 excavation season at Buran-Kaya III; photo © A. Yanevich. b) Anthropological study of human remains in the high containment laboratory at the IJM; photo © E.M. Geigl. c) Images of the human parietal fragment from 6-1.a, *BuranKaya3A*, analyzed in the present study, taken under a stereomicroscope (Leica MZ FLIII, PLANAPO 0.63X); photo © E.M. Geigl.

**Table S1.** Comparative $^{14}$C dates. All the dates were calibrated using the software OxCal v4.3.2 based on the IntCal13 calibration data set[1]. The calibrated dates are rounded to 5.

| Site | Individual | Cultural layer | Reference cultural layer | $^{14}$C (BP) | σ(+) | σ (-) | cal BP (2ρ) | | Laboratory Code | Material dated | Reference datation |
|---|---|---|---|---|---|---|---|---|---|---|---|
| | | | | | | | low | high | | | |
| Peştera cu Oase | Oase1 | No direct cultural association | [2] | 34,290<br>>35,200<br>34,950 | 970<br><br>900 | 870<br><br>890 | <br><br>41,640 | <br><br>37,580 | GrA-22810<br>Ox-A-11711<br>Combined age | Direct date on human remains | [2] |
| Kostenki (Martina Gora) | Kostenki14 | No direct cultural association | [3] | 33,250 | 500 | 500 | 38,680 | 36,260 | OxA-X-2395-15 | Direct date on human remains | [4] |
| **Buran-Kaya III** | **BuranKaya3A** | **Layer 6-1 Gravettian** | [5] | **31,900** | **240** | **220** | **36,310** | **35,240** | **GrA-37938** | **Dated on human remains from the same layer** | [5] |
| Goyet | Goyet Q116-1 | No direct cultural association | [6] | 30,880 | 170 | 160 | 35,160 | 34,430 | GrA-46175 | Direct date on human remains | [6] |
| Sunghir | Sunghir3 | Cultural assignment discussed | For the whole site : Late Sungirian[7,8] Late Streletskian [4,9–11] Eastern Gravettian[12,13] | 30,000<br>24,830 | 550<br>110 | 550<br>110 | 35,150 | 33,030 | OxA-X-2395-7<br>OxA-15754 | Direct date on human remains | [4] |
| Kostenki 12 (Volkovskaya site) | Kostenki12 | Attributed to Gorodtsovian | [14] | 28,500 | 140 | 140 | 32,995 | 31,840 | GrA-5552 | Associated charcoal | [14] |
| Pavlov I | Pavlov1 | Gravettian (evolved Pavlovian) | [15] | 26,170 | 450 | 450 | 31,100 | 29,420 | GrN-20391 | Associated charcoal | [15] |
| Dolní Věstonice II | Vestonice16 | Gravettian (evolved Pavlovian) | [15] | 25,740<br>25,570 | 210<br>280 | 210<br>280 | 30,570<br>30,535 | 29,390<br>29,035 | GrN-15277<br>GrN-15276 | Associated charcoal | [15] |

**Table S2.** Shotgun library results

| Library | Total reads >27bp | hg37d5 Mapped reads | PCR duplicates removed | Mapping Quality 20 | Average length (bp) | % endogenous DNA |
|---|---|---|---|---|---|---|
| SP61-8-17_BKA4_B | 6441522 | 26702 | 25125 | 19480 | 35.0 | 0.41% |
| SP62-1_BKA4_B_UNG | 27957579 | 117303 | 107828 | 85025 | 37.1 | 0.42% |
| SP62-2_BKA4_B | 10456937 | 75619 | 63145 | 46579 | 35.5 | 0.72% |
| SP62-3_BKA4_B | 40315968 | 156287 | 151945 | 117997 | 39.7 | 0.39% |
| SP62-4_BKA4_B | 38353291 | 151652 | 147546 | 115046 | 39.6 | 0.40% |
| SP62-5_BKA4_B | 34800662 | 124997 | 122028 | 95191 | 39.0 | 0.36% |
| SP62-6_BKA4_B | 32251522 | 109869 | 107635 | 84208 | 38.5 | 0.34% |
| SP62-7_BKA4_B | 36758191 | 141531 | 138038 | 107792 | 38.7 | 0.39% |
| SP62-8_BKA4_B | 20324986 | 108640 | 106323 | 82621 | 38.8 | 0.53% |
| Total reads | 247660658 | 1012600 | 969613 | 753939 | 38.0 | 0.44% |

**Table S3.** Y SNP summary with Palaeolithic comparisons

| **Y haplogroup** | **BuranKaya3A SNPs** | | **Palaeolithic haplogroups** |
|---|---|---|---|
| | **Derived** | **Ancestral** | |
| **BT** | **6** | 0 | |
| **CT** | **1** | 0 | Kostenki12[16], 3/5 Natufians[17] |
| **C** | **1** | 0 | |
| C1a* | | | GoyetQ116-1[16] |
| C1a2 | 0 | 7 | Sunghir3[18], Vestonice16[16] |
| C1b* | | | Kostenki14[3] |
| C1b1a1 | 0 | 1 | |
| C1b1a2 | 0 | 6 | |
| C1b1a3 | 0 | 1 | |
| C1b2a | 0 | 1 | |
| C2 | 0 | 7 | |

**Table S4.** Neanderthal content of BuranKaya3A and three ancient genomes recalculated using the BuranKaya3A subset of 6,252 Neanderthal- and modern human-derived SNPs, compared with the previously published median results calculated from either archaic ancestry or *f*4-ratio method from Fu et al., 2016[16].

| | **BK SNP subset** | **95%CI** | **Fu-Ancestry** | **Fu-*f*4** |
|---|---|---|---|---|
| BuranKaya3A | 3.55% | 0.0224 | | |
| Oase | 8.82% | 0.0973 | 7.5% | 9.9% |
| Kostenki14 | 2.94% | 0.0205 | 2.8% | 3.6% |
| Ust-Ishim | 4.69% | 0.0236 | 3.0% | 4.4% |

**Table S5.** Mitochondrial sequence references

| | Haplogroup | Age* | Location | Culture | Reference |
|---|---|---|---|---|---|
| African | L3 | modern | West Africa (Yoruba) | - | AF347014.1 |
| BuranKaya3A | N1 | 35775 | Crimea | Early Gravettian | this study |
| Cioclovina1 | U | 33212 | Romania | - | [6] |
| Fumane2 | R | 39805 | Italy | - | [19] |
| Goyet2878-21 | U5 | 26662 | Belgium | Gravettian | [6] |
| GoyetQ116-1 | M | 34795 | Belgium | Aurignacian | [6] |
| GoyetQ376-19 | U2 | 27515 | Belgium | Gravettian | [6] |
| GoyetQ376-3 | M | 33540 | Belgium | Aurignacian | [6] |
| GoyetQ53-1 | U2 | 27975 | Belgium | Gravettian | [6] |
| GoyetQ55-2 | U2 | 27520 | Belgium | Gravettian | [6] |
| GoyetQ56-16 | U2 | 26320 | Belgium | Gravettian | [6] |
| Kostenki14 | U2 | 37470 | Russia | - | [20] |
| LaRochette | M | 27592 | France | | [6] |
| Muierii1 | U6 | 35204 | Romania | - | [21] |
| N1a | N1a | NA | NA | NA | N1a defining mutations |
| N1b | N1b | NA | NA | NA | N1b defining mutations |
| Natufian9 | N1 | 10800 | Isreal | Natufian | [17] |
| Oase1 | N | 39610 | Romania | - | [22] |
| Paglicci108 | U2'3'4'7'8'9 | 28396 | Italy | Gravettian | [6] |
| Paglicci133 | U8c | 33000 | Italy | Gravettian | [6] |
| Salkhit | N | 34425 | Mongolia | - | [23] |
| Sunghir1 | U8 | 32820 | Russia | Eastern Gravettian/Streletskian | [18] |
| Sunghir2 | U2 | 34230 | Russia | Eastern Gravettian/Streletskian | [18] |
| Sunghir3 | U2 | 34090 | Russia | Eastern Gravettian/Streletskian | [18] |
| Sunghir4 | U2 | 33988 | Russia | Eastern Gravettian/Streletskian | [18] |
| Tianyuan | R | 40328 | China | - | [23] |
| Ust-Ishim | R | 45020 | Siberia | - | [24] |

| | | | | | |
|---|---|---|---|---|---|
| Vestonice13 | U8 | 31155 | Czech Republic | Gravettian | [25] |
| Vestonice14 | U5 | 31155 | Czech Republic | Gravettian | [25] |
| Vestonice16 | U5 | 29980 | Czech Republic | Gravettian | [6] |
| Vestonice43 | U5 | 29977 | Czech Republic | Gravettian | [6] |

*median value (calBP) from dates given in reference

**Table S6.** Genome sequence references

| | Age* | Location | Culture | Reference |
|---|---|---|---|---|
| AfontovaGora3 | 16710 | Russia | - | [16] |
| BerryAuBac | 7245 | France | Mesolithic | [16] |
| Bichon | 13665 | Switzerland | Azilian | [26] |
| Bockstein | 8265 | Germany | Mesolithic | [16] |
| Brillenhohle | 14780 | Germany | Magdelenian | [16] |
| BuranKaya3A | 35775 | Crimea | Gravettian | This study |
| Burkhardtshohle | 14615 | Germany | Magdelenian | [16] |
| Chaudardes1 | 8205 | France | Mesolithic | [16] |
| Cioclovina1 | 32435 | Romania | Mesolithic | [16] |
| Continenza | 10855 | Italy | Mesolithic | [16] |
| ElMiron | 18720 | Spain | Magdelenian | [16] |
| Falkenstein | 9200 | Germany | Mesolithic | [16] |
| GoyetQ116-1 | 34795 | Belgium | Aurignacian | [16] |
| GoyetQ-2 | 15005 | Belgium | Magdelenian | [16] |
| GoyetQ376-19 | 27515 | Belgium | Gravettian | [16] |
| GoyetQ53-1 | 27975 | Belgium | Gravettian | [16] |
| GoyetQ56-16 | 26320 | Belgium | Gravettian | [16] |
| HohleFels49 | 15130 | Germany | Magdelenian | [16] |
| HohleFels79 | 14670 | Germany | Magdelenian | [16] |
| Hotu | 8878 | Iran | Mesolithic | [17] |
| Iboussieres39 | 11725 | France | Epipaleolithic | [16] |
| Karelia | 8375 | Russia | Mesolithic | [27] |
| Kostenki12 | 32415 | Russia | Attributed to Gorodtsovian | [16] |
| Kostenki14 | 37470 | Russia | - | [16,28] |
| Kotias | 9720 | Georgia | Mesolithic | [26] |
| KremsWA3 | 30970 | Austria | Gravettian | [16] |

| | | | | |
|---|---|---|---|---|
| LaBrana1 | 7815 | Spain | Mesolithic | [29] |
| LesCloseaux13 | 9562 | France | Mesolithic | [16] |
| Loschbour | 8050 | Luxembourg | Mesolithic | [30] |
| Mal'ta | 24305 | Russia | Mal'ta - Buret' | [31] |
| Muierii2 | 33300 | Romania | - | [16] |
| Natufian9 | 10800 | Israel | Natufian | [17] |
| Natufian6 | 10800 | Israel | Natufian | [17] |
| Oase1 | 39610 | Romania | - | [22] |
| Ofnet | 8245 | Germany | Mesolithic | [16] |
| Ostuni1 | 27620 | Italy | Gravettian | [16] |
| Ostuni2 | 28975 | Italy | Gravettian | [16] |
| Paglicci108 | 28396 | Italy | Gravettian | [16] |
| Paglicci133 | 32895 | Italy | Gravettian | [16] |
| Pavlov1 | 30260 | Czech Republic | Gravettian | [16] |
| Ranchot88 | 10085 | France | Mesolithic | [16] |
| Rigney1 | 15465 | France | Magdelenian | [16] |
| Rochedane | 12960 | France | Epipaleolithic | [16] |
| Satsurblia | 13255 | Georgia | Epigravettian | [26] |
| Sunghir3 | 34090 | Russia | Eastern Gravettian/Streletskian | [18] |
| Tianyuan | 40328 | China | - | [32] |
| Ust-Ishim | 45020 | Siberia | - | [24] |
| Vestonice13 | 31155 | Czech Republic | Gravettian | [16] |
| Vestonice14 | 31155 | Czech Republic | Gravettian | [16] |
| Vestonice15 | 31155 | Czech Republic | Gravettian | [16] |
| Vestonice16 | 29980 | Czech Republic | Gravettian | [16] |
| Vestonice43 | 29977 | Czech Republic | Gravettian | [16] |
| Villabruna | 13980 | Italy | Epigravettian | [16] |

*median value (calBP) from dates given in reference

**Table S7.** *f3*(BuranKaya3A, *x*, Mbuti) results

| | | | All SNPS | | | | Transversions only | | | |
|---|---|---|---|---|---|---|---|---|---|---|
| | | | **f3** | **sterr** | **Z** | **SNPs** | **f3** | **sterr** | **Z** | **SNPs** |
| BuranKaya3A | AfontovaGora3 | Mbuti | 0.384319 | 0.035709 | 10.762 | 1327 | 0.350285 | 0.07188 | 4.873 | 298 |
| BuranKaya3A | BerryAuBac | Mbuti | 0.332715 | 0.079569 | 4.181 | 226 | 0.341718 | 0.144269 | 2.369 | 52 |
| BuranKaya3A | Bichon | Mbuti | 0.402716 | 0.015075 | 26.714 | 8262 | 0.413134 | 0.021643 | 19.088 | 3862 |
| BuranKaya3A | Bockstein | Mbuti | 0.419447 | 0.147753 | 2.839 | 83 | 0.528223 | 0.459984 | 1.148 | 11 |
| BuranKaya3A | Brillenhohle | Mbuti | 0.128968 | 0.103485 | 1.246 | 51 | 0.091156 | 0.174642 | 0.522 | 10 |
| BuranKaya3A | Burkhardtshohle | Mbuti | 0.385832 | 0.117609 | 3.281 | 131 | 0.611612 | 0.338867 | 1.805 | 28 |
| BuranKaya3A | Chaudardes1 | Mbuti | 0.515568 | 0.083193 | 6.197 | 309 | 0.424476 | 0.173462 | 2.447 | 65 |
| BuranKaya3A | Cioclovina1 | Mbuti | 0.520214 | 0.187446 | 2.775 | 62 | 0.175386 | 0.345346 | 0.508 | 15 |
| BuranKaya3A | Continenza | Mbuti | 0.477686 | 0.263526 | 1.813 | 48 | 0.592877 | 0.81353 | 0.729 | 13 |
| BuranKaya3A | ElMiron | Mbuti | 0.391415 | 0.021648 | 18.081 | 3547 | 0.354761 | 0.035589 | 9.968 | 1066 |
| BuranKaya3A | Falkenstein | Mbuti | 0.348055 | 0.074946 | 4.644 | 231 | 0.401859 | 0.213542 | 1.882 | 37 |
| BuranKaya3A | GoyetQ116-1 | Mbuti | 0.402532 | 0.022416 | 17.958 | 3659 | 0.387871 | 0.045382 | 8.547 | 795 |
| BuranKaya3A | GoyetQ-2 | Mbuti | 0.453124 | 0.077433 | 5.852 | 263 | 0.605961 | 0.191084 | 3.171 | 61 |
| BuranKaya3A | GoyetQ376-19 | Mbuti | 0.276116 | 0.106582 | 2.591 | 88 | 0.223004 | 0.136924 | 1.629 | 17 |
| BuranKaya3A | GoyetQ53-1 | Mbuti | 0.185602 | 0.175285 | 1.059 | 34 | 0.323704 | 0.534602 | 0.606 | 6 |
| BuranKaya3A | GoyetQ56-16 | Mbuti | 0.553546 | 0.272756 | 2.029 | 28 | 0.352032 | 0.56187 | 0.627 | 7 |
| BuranKaya3A | HohleFels49 | Mbuti | 0.543713 | 0.09323 | 5.832 | 262 | 0.494036 | 0.250399 | 1.973 | 41 |
| BuranKaya3A | HohleFels79 | Mbuti | 0.240254 | 0.153623 | 1.564 | 42 | 0.091954 | 0.021719 | 4.234 | 11 |
| BuranKaya3A | Hotu | Mbuti | 0.376847 | 0.050784 | 7.421 | 691 | 0.261641 | 0.088716 | 2.949 | 153 |
| BuranKaya3A | Iboussieres39 | Mbuti | 0.504506 | 0.331982 | 1.52 | 29 | 0.247283 | 0.397081 | 0.623 | 8 |
| BuranKaya3A | Karelia | Mbuti | 0.422672 | 0.01737 | 24.333 | 7067 | 0.432769 | 0.024782 | 17.463 | 3247 |
| BuranKaya3A | Kostenki12 | Mbuti | 0.406626 | 0.075608 | 5.378 | 310 | 0.489396 | 0.174374 | 2.807 | 88 |
| BuranKaya3A | Kostenki14 | Mbuti | 0.430879 | 0.01726 | 24.964 | 7311 | 0.427491 | 0.024095 | 17.742 | 3089 |
| BuranKaya3A | Kotias | Mbuti | 0.393417 | 0.014959 | 26.3 | 8252 | 0.397889 | 0.021343 | 18.643 | 3869 |
| BuranKaya3A | KremsWA3 | Mbuti | 0.386445 | 0.039692 | 9.736 | 1031 | 0.527171 | 0.098493 | 5.352 | 231 |

| | | | | | | | | | | |
|---|---|---|---|---|---|---|---|---|---|---|
| BuranKaya3A | LaBrana1 | Mbuti | 0.405692 | 0.015797 | 25.681 | 7271 | 0.414868 | 0.022658 | 18.31 | 3305 |
| BuranKaya3A | LesCloseaux13 | Mbuti | 0.156012 | 0.104236 | 1.497 | 35 | 0.156353 | 0.200758 | 0.779 | 5 |
| BuranKaya3A | Loschbour | Mbuti | 0.420236 | 0.014939 | 28.13 | 8324 | 0.425356 | 0.021072 | 20.186 | 3900 |
| BuranKaya3A | Mal’ta | Mbuti | 0.395182 | 0.017644 | 22.398 | 6446 | 0.402512 | 0.025462 | 15.808 | 2966 |
| BuranKaya3A | Muierii | Mbuti | 0.481571 | 0.067999 | 7.082 | 477 | 0.68802 | 0.142958 | 4.813 | 142 |
| BuranKaya3A | Natuian9 | Mbuti | 0.374678 | 0.027978 | 13.392 | 2070 | 0.402556 | 0.061442 | 6.552 | 454 |
| BuranKaya3A | Natufian6 | Mbuti | 0.293264 | 0.049968 | 5.869 | 617 | 0.447189 | 0.122606 | 3.647 | 145 |
| BuranKaya3A | Oase1 | Mbuti | 0.329862 | 0.035003 | 9.424 | 1158 | 0.301321 | 0.055523 | 5.427 | 459 |
| BuranKaya3A | Ofnet | Mbuti | 0.397787 | 0.262171 | 1.517 | 23 | 0.150049 | 0.314613 | 0.477 | 5 |
| BuranKaya3A | Ostuni1 | Mbuti | 0.365527 | 0.03162 | 11.56 | 1622 | 0.418664 | 0.060023 | 6.975 | 418 |
| BuranKaya3A | Ostuni2 | Mbuti | 0.45448 | 0.165329 | 2.749 | 83 | 0.307497 | 0.521756 | 0.589 | 15 |
| BuranKaya3A | Paglicci108 | Mbuti | 0.773895 | 0.509567 | 1.519 | 13 | -0.125 | 1.608895 | -0.078 | 5 |
| BuranKaya3A | Paglicci133 | Mbuti | 0.322502 | 0.069246 | 4.657 | 319 | 0.402489 | 0.164335 | 2.449 | 68 |
| BuranKaya3A | Pavlov1 | Mbuti | 0.449836 | 0.080795 | 5.568 | 299 | 0.478315 | 0.193927 | 2.466 | 65 |
| BuranKaya3A | Ranchot88 | Mbuti | 0.398034 | 0.032273 | 12.333 | 1738 | 0.387009 | 0.064378 | 6.011 | 343 |
| BuranKaya3A | Rigney1 | Mbuti | 0.41524 | 0.103769 | 4.002 | 150 | 0.591176 | 0.454989 | 1.299 | 27 |
| BuranKaya3A | Rochedane | Mbuti | 0.380682 | 0.039607 | 9.612 | 995 | 0.515008 | 0.091489 | 5.629 | 215 |
| BuranKaya3A | Satsurblia | Mbuti | 0.408289 | 0.017349 | 23.534 | 5702 | 0.42183 | 0.024122 | 17.487 | 2711 |
| BuranKaya3A | Sunghir3 | Mbuti | 0.448631 | 0.015876 | 28.258 | 8348 | 0.448689 | 0.022101 | 20.302 | 3922 |
| BuranKaya3A | Tianyuan | Mbuti | 0.365052 | 0.015869 | 23.004 | 5690 | 0.364851 | 0.023197 | 15.728 | 2558 |
| BuranKaya3A | Ust-Ishim | Mbuti | 0.401416 | 0.013894 | 28.891 | 9358 | 0.414576 | 0.019685 | 21.06 | 4375 |
| BuranKaya3A | Vestonice13 | Mbuti | 0.394869 | 0.047024 | 8.397 | 679 | 0.297689 | 0.080165 | 3.713 | 189 |
| BuranKaya3A | Vestonice14 | Mbuti | 0.141359 | 0.216404 | 0.653 | 20 | 0.335942 | 0.457878 | 0.734 | 5 |
| BuranKaya3A | Vestonice15 | Mbuti | 0.406163 | 0.119414 | 3.401 | 148 | 0.167227 | 0.193605 | 0.864 | 30 |
| BuranKaya3A | Vestonice16 | Mbuti | 0.435233 | 0.0218 | 19.965 | 4088 | 0.437643 | 0.035735 | 12.247 | 1326 |
| BuranKaya3A | Vestonice43 | Mbuti | 0.373642 | 0.048209 | 7.751 | 745 | 0.419048 | 0.08648 | 4.846 | 253 |
| BuranKaya3A | Villabruna | Mbuti | 0.377008 | 0.017081 | 22.072 | 5295 | 0.349925 | 0.026086 | 13.414 | 1852 |
